# Detecting Cd adaptation footprint in C. riparius with a multi-genomic approach

**DOI:** 10.1101/2021.10.13.464202

**Authors:** Halina Binde Doria, Pauline Hannapel, Markus Pfenninger

## Abstract

Evolutionary processes and acquired tolerance to toxicants are important factors governing how animals respond to chemical exposure. Evidence for increased tolerance to cadmium (Cd), a widely distributed toxic metal in aquatic environments, in *Chironomus riparius* is conflicting and still questioned if it happens through phenotypic plasticity or genetic adaptation. The present study considered the relevance of directional environmental changes by increasing contaminant concentration in a multigenerational selection experiment. Evaluation of measurable life-cycle traits, transcriptomic responses and quantitative genetics from an evolve and resequencing (E&R) experiment were integrated to assess the potential of *C. riparius* to adapt to Cd. Survival tests revealed some adaptation to Cd exposure. Genomic analyses showed a strong, genome-wide selective response in all replicates, emphasizing that even control laboratory conditions continually exert selective pressure. The integration of transcriptomic and genomic data could isolate the genes related to Cd acquired resistance. Those genes could be linked to an efflux of metals. Therefore, it is possible to conclude that *C. riparius* can endure long-term Cd exposure also through genetic adaptation.

## 1. INTRODUCTION

Natural populations are typically chronically exposed to pollutants over the course of many generations and therefore it is important to study the long-term responses to chemical exposure (Beaudouin et al., 2012; Im et al., 2019; Klerks and Weis, 1987). Long term exposure may result in evolution of resistance and reduced genetic diversity (Klerks et al., 2011; Nowak et al., 2009). Both may impact the ability of a population to cope with subsequent changes in stress conditions and therefore can have important implications regarding ecotoxicological risk assessments and long term survival of the species (Becker and Liess, 2015; Chaumot et al., 2009; Zhang et al., 2019).

Consequently, it is crucial to add an evolutionary perspective to understand and quantify evolutionary dynamics of chronically exposed populations (Beaudouin et al., 2012; Becker and Liess, 2015; Straub et al., 2020). But despite the increasing number of multigenerational investigations, most are focused on observed phenotypical changes of measured life-traits (Doria and Pfenninger, 2021; Rodríguez-Romero et al., 2021). However, phenotypes may be plastic and, although influenced by current and historical environmental conditions, they can be reversible changes (Shaw et al., 2019). Thus, commonly measured life-history traits are fitness components that tend to possess low heritability (Mousseau and Roff, 1987). Additionally, although high-throughput “omics” technologies are facilitating the detection of new biomarkers and pathway responses to a toxicant (Canzler et al., 2020; Ebner, 2021), there is a remarkable non utilization of the evolve and resequence (E&R) approach in ecotoxicology. This last would ultimately enable molecular evolution to be monitored in real time on a genome-wide scale and open up the possibility to identify the loci contributing to genetic adaptation (Kofler and Schlötterer, 2014; Long et al., 2015).

*Chironomus riparius*, known as harlequin-fly, is a non-biting midge and during its larval stage is a sediment-dwelling organism (Horváth et al., 2011). Its larvae are therefore widely employed in sediment-water toxicity tests and currently the species is used in two internationally recognized standardized tests (OECD, 2011, 2010). Further, due to its short life-cycle (Doria and Pfenninger, 2021; Foucault et al., 2019) and ability to rapid adapt to environmental variables and pollution (Beaudouin et al., 2012; Pfenninger and Foucault, 2020a, 2020b), it is an emerging model organism for E&R studies.

Cadmium (Cd) is both a legacy contaminant and a present-day pollutant of great concern (Dong et al., 2021; Shaw et al., 2019). It can promote oxidative stress and negatively affect immune, inflammatory and reproductive endpoints, being one of the most toxic metals present in aquatic environments (Blickley et al., 2014; Doria and Pfenninger, 2021; Martín-Folgar and Martínez-Guitarte, 2019). But although, it is listed as a carcinogenic and mutagenic substance (Wang et al., 2019), it was recently demonstrated that it does not affect germ cell mutation rates in *C. riparius* (Doria et al., 2021)

Cd accumulation dynamics by living organisms is rather determined by its bioavailability than the total concentration in soils and sediments (Dong et al., 2021; Pauget et al., 2013). Usually, contamination sources may modulate Cd bioavailability and the newly introduced Cd through recent anthropogenic activities has higher bioavailability (Dong et al., 2021; Pauget et al., 2013).

The heritability of acquired Cd resistance has distinct reports for numerous different organisms (Dallinger and Höckner, 2013). In the case of *C. riparius*, early studies attributed a strong heritable genetic background to it (J.F. Postma et al., 1995; Jaap F Postma et al., 1995; Postma and Davids, 1995). However, more recent studies claimed that a decreased sensitivity to Cd, regardless of the exposure scenario, does not always occur, but when it happens, it relied rather on plasticity and was only weakly heritable (Doria and Pfenninger, 2021; Marinković et al., 2012; Vogt et al., 2010).

These seemingly contradictory results might be due to the genetic background of the experimentally exposed populations, since available genetic variation is a crucial factor for evolutionary adaptation (Barrett and Schluter, 2008; Dallinger and Höckner, 2013). However, the strength and duration of selection also play a big role (Charlesworth, 2009). A directional selection of favourable genotypes (given respective standing genetic variation within the population), incurs that the impact must be profound and constant as to represent a strong selective force (Charlesworth, 2009; Dallinger and Höckner, 2013; Ward and Robinson, 2005). The normal scenario in most bioassays is to impose a sudden large change in the environment, after which the environment is kept constant (Gorter et al., 2017). Yet, organisms in natural environments typically endure gradually changing conditions. Therefore, an overly strong selection may bias the outcome of evolutionary experiments because of the loss of genetic variation resulting from the demographic crash following an instantly strong environmental change (Klerks et al., 2011). As a result, laboratory experiments may give a biased impression on the evolutionary potential in the field.

In the present study, we made use of gradually rising Cd concentrations in cultures over eight generations to simulate environmentally realistic conditions. Further, we employed both methods commonly used to study experimentally the evolution of resistance: (i) Time-series assessments of fitness components to evaluate whether contaminant-exposed replica showed increased Cd resistance in comparison to negative control populations. (ii) An E&R approach to infer the traces of genome-wide molecular selection imposed by the selective regime applied. In addition, we used a multi-genomic data integration, namely and whole RNA and DNA sequencing.

With those tools we aimed to answer if a tolerance acquisition to Cd occurred, was it then heritable and thus induce genetic adaptation of the population? We also inferred the effect on regulation of the transcription level and the changes in allele frequencies.

## 2. MATERIAL AND METHODS

### 2.1. Source population and establishment of the test replicates

The *C. riparius* population used in the study originated from a native population (Hasselbach, Hessen, Germany; 50.167562°N, 9.083542°E) and has been maintained for several generations as a large in-house laboratory culture. This population has been previously used in a handful of ecotoxicology and ecology studies (Carrasco-Navarro et al., 2021; Doria and Pfenninger, 2021; Pfenninger and Foucault, 2020b).

Twenty egg ropes from the previously described population were collected and, after successful hatching, they were pooled together to start six different replicates. At least 1000 newly hatched larvae started the population of each replicate. Three replicates were assigned to control conditions and the other three were treated with increasing Cd concentration. Which meant that, for the Cd exposed groups, at every twenty-eight days - around the length of one generation - a nominal concentration of 50 ug/L of Cd were added to the replicates. All replicates were maintained under 23°C, 60% humidity and photoperiod of 16:8 light:dark, following the OECD guideline 233 (OECD, 2010). The culturing conditions for all the replicates followed the method described in Foucault *et al*. (2019).

Replicates were raised at these conditions for eight generations. After the sixth generation six egg ropes from the Cd exposed replicates - two from each replicate – were collected and pooled after hatching. From this pool, 1000 larvae were sorted and put in control conditions to produce the recovery group, which was maintained for further two generations.

### 2.2. Test compound and chemical analysis

A stock solution of cadmium chloride salt (Merck, Germany) of 10 mg/ L was used. The salt was diluted in deionized dechlorinate water and kept under 4 °C. Cd concentrations in water and sediment were measured at each fifty days, around the length of two generations, in the replicates and at the end of the life cycle experiments. Cd concentration in water samples were determined by inductively coupled plasma mass spectrometry (detection limit 0.0002 mg/L) (DIN EN ISO 17294e2; 2014). Cd concentrations in the sediment was assessed after extraction in a HCl + HNO_3_ mixture (aqua regia) (DIN EN 13657; 2003) and analysed by inductively coupled plasma optical emission spectrometry (detection limit of 0.2 mg/kg) (DIN EN ISO 11885; 2009).

### 2.3. Acute and chronic survival experiments

Common garden experiments were used to perform the survival tests, which took place after the first and eighth generation of the Cd pre-treated and control replicates and after the second generation of recovery group. These three different time points were used to investigate possible acclimation or adaptation during the increasing Cd exposure. At each time point, fifteen egg clutches from each pre-treated replicate and the recovery group were collected and allowed to hatch. Ten fully hatched egg clutches, five in the case of the recovery group. were then randomly chosen for the tests. Individuals coming from the same egg clutch are referred to as families (Figure 1).

**Figure 1:**
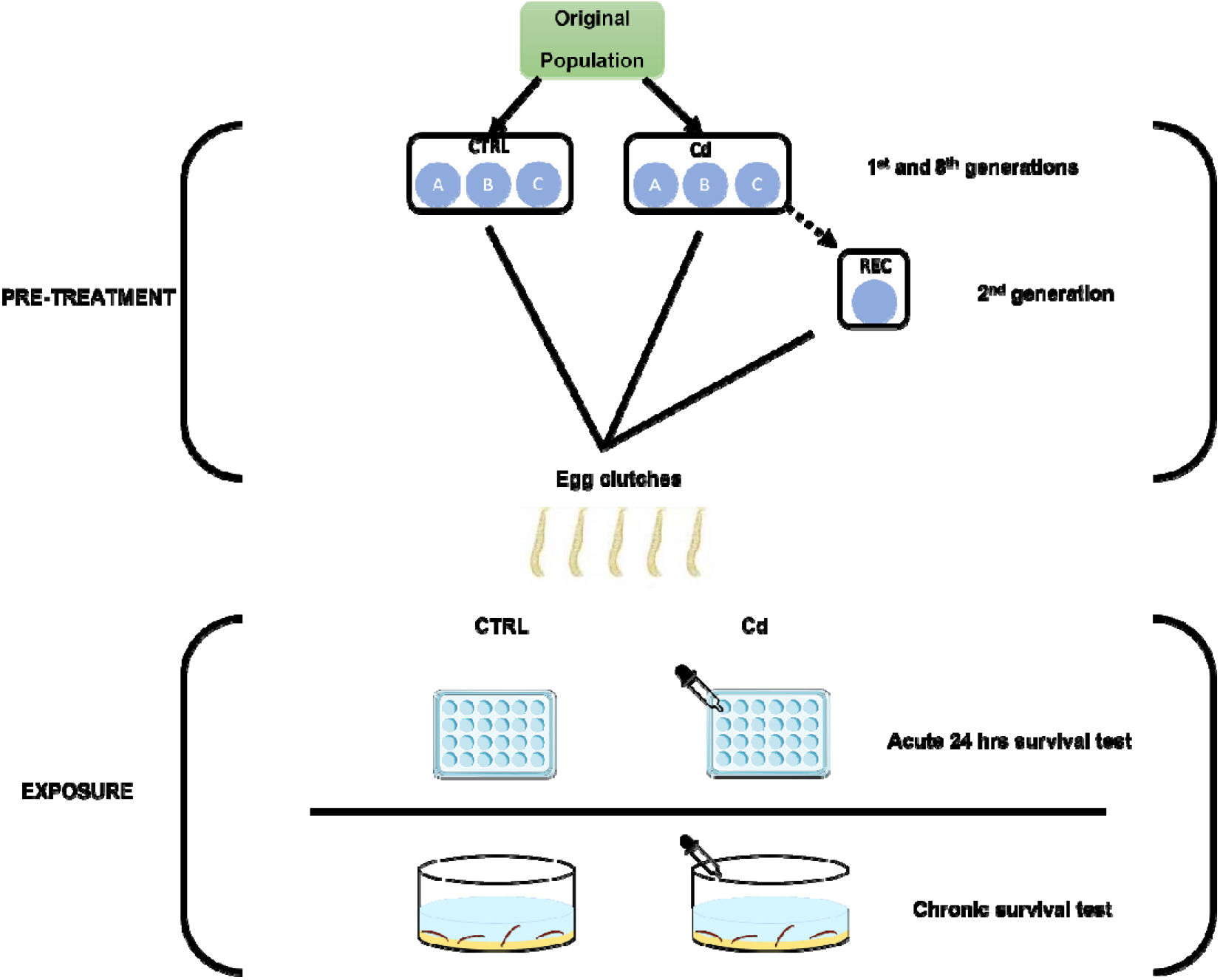
Schema of the experimental design used to record acute and chronic mortality rate using a common garden approach.

The acute examination consisted of twenty 24-well plates per test group, of which Cd was added to ten of these plates and ten remained untreated. The recovery group consisted of ten 24-well plates per test group, of which Cd was added to five of these plates and five remained untreated Therefore, each well was filled with 2 mL of medium or 2 mL of medium + 50 ug/L of Cd. At each well one newly hatched larva was introduced and raised without feeding to avoid medium alteration. Plates were covered and mortality was assessed after 24 h. Each test group consisted of 360 larvae (24 larvae × 5 families × 3 population replicates), except from the recovery group that consisted in 120 larvae (24 larvae × 5 families). The aim of the acute test was to find out whether the chironomids were already, in the first larval stage, negatively impaired by the Cd exposure.

The chronic survival test was performed according to Foucault *et al*. (2018). Each group evaluated consisted of twenty replicates in generation eight, from which ten were spiked with a nominal concentration of 50 ug/L of Cd in the water. For the first generation and the recovery assessments, each group evaluated consisted of ten replicates, from which five were spiked with a nominal concentration of 50 ug/L of Cd in the water. Thirty larvae from each family were added to each glass bowl (Ø20 × 10 cm) with 1,5 cm of sediment and 1.250 L of medium. Test vessels were kept under the same conditions as the replicates culture, i.e. climate room at a temperature of 23 ± 1 °C, 60% relative humidity, light:dark rhythm of 16:8 h photoperiod and were constantly aerated. Water evaporation in the test vessels was compensated for by adding demineralized water. Larvae were fed daily with dried fish food (TetraMin^®^ flakes) according to a feeding schedule (Supplementary Information - S1). Test duration was set at maximum of 28 days. The mortality of pupation and emergence were calculated as the number of individuals not emerged per replicate, median emergence time (EmT50) was calculated as the time when 50% of the female individuals emerged.

To test differences between the three pre-treatments Cd exposed (Cd), control (CTRL) and recovery (REC), a Kruskall-Wallis followed by a post-hoc Dunn’s test for multiple comparisons of independent samples were conducted. Expressed trends of a group between generations were tested with a Bayesian estimation alternative to the t-test (Bååth, 2014; Kruschke, 2013). For graphical representation pre-treatment is showed before (no-Cd or pre-Cd) and separated by a colon from the exposure condition (CTRL or Cd), that way a total of four groups (no-Cd:CTRL, no-Cd:Cd, pre-Cd:CTRL and pre-Cd:Cd) are graphically showed.

For comparison between groups within the same generation data was tested for normality with Shapiro-Wilk test and homogeneity of variances with Levene’s test. Following, one-way analysis of variances (ANOVA) followed by Tukey’s post hoc test was employed. Individual comparisons between Cd exposure and control condition on the recovery group (REC) within each generation was conducted with a two tailed t-test. All the results from t-test, ANOVA and p-values from Tuckey’s post hoc test can be found in the Supplementary Information -S2.

All statistical analysis were done with the software R (Version 4.0.0) using the packages BayesianFirstAid for bayesian estimation alternative to the t-test, dplyr for t-tests and ANOVAs, and the package car for post hoc tests.

### 2.4. Time-series analysis of DNA Poolseq data

To investigate if the replicates could rapidly adapt to the Cd pre-treatment, we performed a whole genome pool-sequencing time-series analysis to track changes in allele frequencies. To this aim, after each 30 days a pool of seventy L4 larvae per replicate were collected. This resulted on a total of forty-eight pools over approximately eight generations, twenty-four from the Cd replicates and twenty-four for the control replicates. DNA was extracted from larval head capsules using the Quiagen blood and tissue extraction kit according to manufacturer’s instructions. Library preparation and sequencing of 150 bp paired end reads using Illumina NovaSeq platform to an expected coverage of 80x were performed by Novogene Europe – UK.

Reads were adapter and quality trimmed using Trimmomatic (Bolger et al., 2014). The clean reads were then mapped against the latest version for *C. riparius* reference genome (Schmidt et al., 2020) using the BWA mem algorithm (Li and Durbin, 2009). Low quality reads were filtered and single nucleotide polymorphisms (SNPs) were initially called using SAMtools (Li et al., 2009). The pipelines Popoolation and Popoolation2 (Kofler et al., 2011a, 2011b) were used to call SNPs, remove indels, and to estimate genetic diversity as Watterson’s theta in the first and last generations of each replicate. Initial and final genetic diversity calculated by Watterson’s theta were tested with paired samples t-test for control and Cd treatment individually.

Allele frequencies for all SNPs with minimal minor allele count above 3 and coverage between 15× and 120× were estimated with the R package PoolSeq (Taus et al., 2017). Random genetic drift was separately simulated for each replicate. First the effective population size (Ne) for each replicate, for intermediate and most evolved generations, were estimated with the JR II plan (Jorde and Ryman, 2007), then expected genetic drift was calculated from the observed allele frequency changes (AFCs) between the first and last generations. SNPs with frequency changes higher than expected under the drift could be identified by chi-square test followed by the Benjamini–Hochberg (BH) correction, to control for false discovery rate, comparing the most evolved to the founder population. A quantile of 99.90% was set to represent the threshold for highly significant SNPs according to chi-square test and used for comparison between real observation and expected drift. Finally, to detect consistent changes across replicates and, therefore, to identify reliable candidates for selected SNPs, a Cochran-Mantel-Haenszel’s (CMH) test was performed merging the three replicates of each experimental group. After, a BH correction for false discovery rate and a cut-off at p < 0.01 was applied (Pfenninger et al., 2021) to generate a list of significant SNPs for each treatment. All calculations and simulations were performed with the R-library PoolSeq (Taus et al., 2017).

Next, highly correlated significant SNPs were merged together, known as hitchhiker SNPs, into haplotype blocks, reducing therefore thousands of candidates to few selected regions (Pfenninger and Foucault, 2020b). Therefore, haplotype selection was done by identifying spatially coherent regions with elevated number of significant SNPs in the filtered output from the CMH test in 10 kb windows with a step size of 500 bp along the scaffolds. Considered regions showed at least one SNP per 10 kb window. Subsequently, to identify the potential target of selection, the SNP with the highest p-value in each previously identified haplotype was filtered. Finally, the annotated genes closest to those selected SNPs were identified as putative selection targets. This generated two lists with the gene’s names, one for CTRL and one for Cd. The duplicates from both lists were removed to yield two list of genes responding uniquely to each treatment. The lists with unique genes responding to each treatment were tested with a Fisher’s exact test for over-representation of GO terms in the category “biological function” with the R-library TopGO (Alexa and Rahnenfuhrer, 2016) using the *weight01* algorithm. Haplotype investigation was done with custom-made phyton scripts following the pipeline already published in Pfenninger & Foucault (2020b).

### 2.5. Gene expression analysis

To investigate the underlying effects of the pre-treatment to Cd at the transcription level we performed a total RNA sequencing analysis. To this aim, after the eighth generation five replicates per group were set following the above-described conditions for the chronic survival test. The experiment used a total of ten egg ropes, five egg ropes were randomly chosen from the Cd pre-treated and the other five originated from the control replicates. From each egg rope 60 larvae were draw, 30 were exposed to 50 ug/L of Cd and 30 were kept under control conditions (0 ug/L of Cd). This resulted on five technical replicates per evaluated group. Name designation of the groups is as previously described for acute and chronic survival experiments.

Six L4 larvae from each replicate were then collected and pooled together two days before emergence, which for the Cd exposed groups happened two days after than the control groups. The pools were immediately homogenized in TRIzol (Invitrogen, ThermoFisher Scientific, Waltham MA, USA) with a pestle and stored at −80°C. RNA extraction of five pools per group was performed with Direct-zol RNA Miniprep (Zymo Research). RNA samples were stored at −80°C after extraction. Single-stranded messenger RNAs (mRNAs) enrichment and conversion to complementary DNA (cDNA), library preparation and sequencing of 150 bp paired end reads using Illumina NovaSeq platform were performed by Novogene Europe – UK.

The quality of the raw reads were assessed using FastQC (Andrews, 2010) and trimming of Illumina adapters was perfomed with Trimmomatic (Bolger et al., 2014). The cleaned reads were then mapped against the latest version of the *C. riparius* genome (NCBI: ERR2696325) (Schmidt et al., 2020) using HISAT2 (Kim et al., 2019). Resulting sam files were converted to bam files and sorted by name with SAMtools (Li et al., 2009). Read count abundances were obtained using HTSeq (Anders et al., 2015), setting the following options: --type mRNA; --mode union; -r name.

To test the individual effects of Cd pre-treatment in gene expression and to track the differences between control conditions and chronic Cd exposure in *C. riparius*, we used the R package Deseq2 v. 1.16.1 (Love et al., 2014) to obtained a list of differentially expressed genes (DEGs) for both conditions. Further, to test the significance of the interaction between pre-treatment and exposure we used the likelihood-ratio (LRT). For the final gene enrichment analysis, the resulting DEG lists were tested with a Fisher’s exact test for over-representation of GO terms in the category “biological function”, representing larger processes completed by several molecular activities, with the R-library TopGO (Alexa and Rahnenfuhrer, 2016) using the *weight01* algorithm.

As a last analysis, to integrate results from the transcriptome to the DNA series analysis, all DEGs resulting from the Cd pre-treatment were tested for overlapping against the significant SNPs list for Cd pre-treatment. The output was then filtered to generate a list of genes present only on the previous list responding uniquely to the Cd pre-treatment. This final list with the overlapping results from DNA time-series analysis and the DEGs found on Cd pre-treatment was equally tested for over-representation of GO terms in the category “biological function” as described above.

For data visualization, the package EnhancedVolcano (Blighe K, Rana S, 2021) was used to generate a volcano plot with the DEGs found on the Cd pre-treatment. The list of DEGs was exported from the DESeq2 analysis results and their counts, log2fold changes and adjusted p-values together with their respective annotations were merged. Package pheatmap (Kolde, 2019) was used to visualize the expression of genes across the replicates and the pcaExplorer (Marini and Binder, 2019) was employed to generate a visualizing the overall effect of experimental covariates between the groups.

## 3. RESULTS

### 3.1. Chemical analysis

The Cd concentration during the chronic life cycle tests conducted on the first, seventh and eighth generations, as well as the first and second generations of the REC group, were below the detection limit (< 0.2 mg/kg of Cd) for sediment samples and 4.1±1.83 ug/L for underlying water samples.

The values for increasing Cd concentrations on the Cd-exposed replicates on sediment started around 0.3 mg/kg of Cd, which is the current LOEC for Cd exposure on *C. riparius* (Doria and Pfenninger, 2021), until about 1.3 mg/kg of Cd. Reversely, the water samples showed higher concentrations on the second and fourth generations varying from 0.7 to 1.2 ug/L that decreased to about 0.5 ug/L on the last measured generation (Table 1).

**Table 1:** Cd concentrations on the overlying water and sediments from the second until the eighth generation measured on the replicates during the evolution experiment.

| replicate/<br>generation | water (ug/L) |  |  |  | sediment (mg/kg) |  |  |  |
| --- | --- | --- | --- | --- | --- | --- | --- | --- |
|  | 2 | 4 | 6 | 8 | 2 | 4 | 6 | 8 |
| **A** | 1.2 | 1.1 | $<0.2$ | 0.4 | 0.3 | 0.3 | 0.4 | 1.3 |
| **B** | 1.2 | 0.7 | $<0.2$ | 0.4 | 0.4 | 0.4 | 0.7 | 1.5 |
| **C** | 0.7 | 1.2 | 0.6 | 0.6 | $<0.2$ | 0.3 | 0.3 | 1 |

### 3.2. Acute and chronic life-cycle experiments

The three pre-treatments Cd, CTRL and REC showed that REC was not different from either Cd exposed or control replicates (Cd x REC p-value = 0.1944; CTRL x REC p-value = 0.3169), but the pre-treatments Cd and CTRL were significantly different (p-value = 0.0034).

The Bayesian approach identified that there was an increasing trend, compromised between 12-19%, of larvae survival in the acute mortality test between the first and last evaluated generations when pre-treated with Cd. Both Cd pre-treated groups independent from exposure scenario showed that the difference of the means have more than 99% probability of being less than zero (difference of the means = −12 [−18, −7] and −19 [−25, −12]; probability of difference of the means = 99.7% < 0 < 0.3% and 99.9% < 0 < 0.1%, for pre-Cd:CTRL and pre-Cd:Cd groups, respectively). Overall, all the groups in all evaluated generations demonstrated survival higher than 80% (Figure 2a). Moreover, a slight rising trend in chronic survival, represented by the number of emerged adults, was detected in the pre-Cd:Cd, with survival improvement around 17% (difference of the means = −17 [−40, 6.2]; probability of difference of the means = 93.7% < 0 < 6.3%) (Figure 2b). EmT50 trend, the day at which 50% emergence occurred, showed a trend of slightly reduced time between the first and the most evolved generation in the no-Cd:CTRL group (difference of the means = 0.96 [−1, 2.8]; probability of difference of the means = 88.4% < 0 < 11.6%). The opposite happened for the no-Cd:Cd group, in which the EmT50 showed an increasing trend (difference of the means = −1.2 [−3.7, 1.5]; probability of difference of the means = 85.1% < 0 < 14.9%). Groups exposed to Cd, independent from the pre-exposure scenario and generation, showed higher EmT50, extending their development in at least two days (Figure 2c).

**Figure 2:**
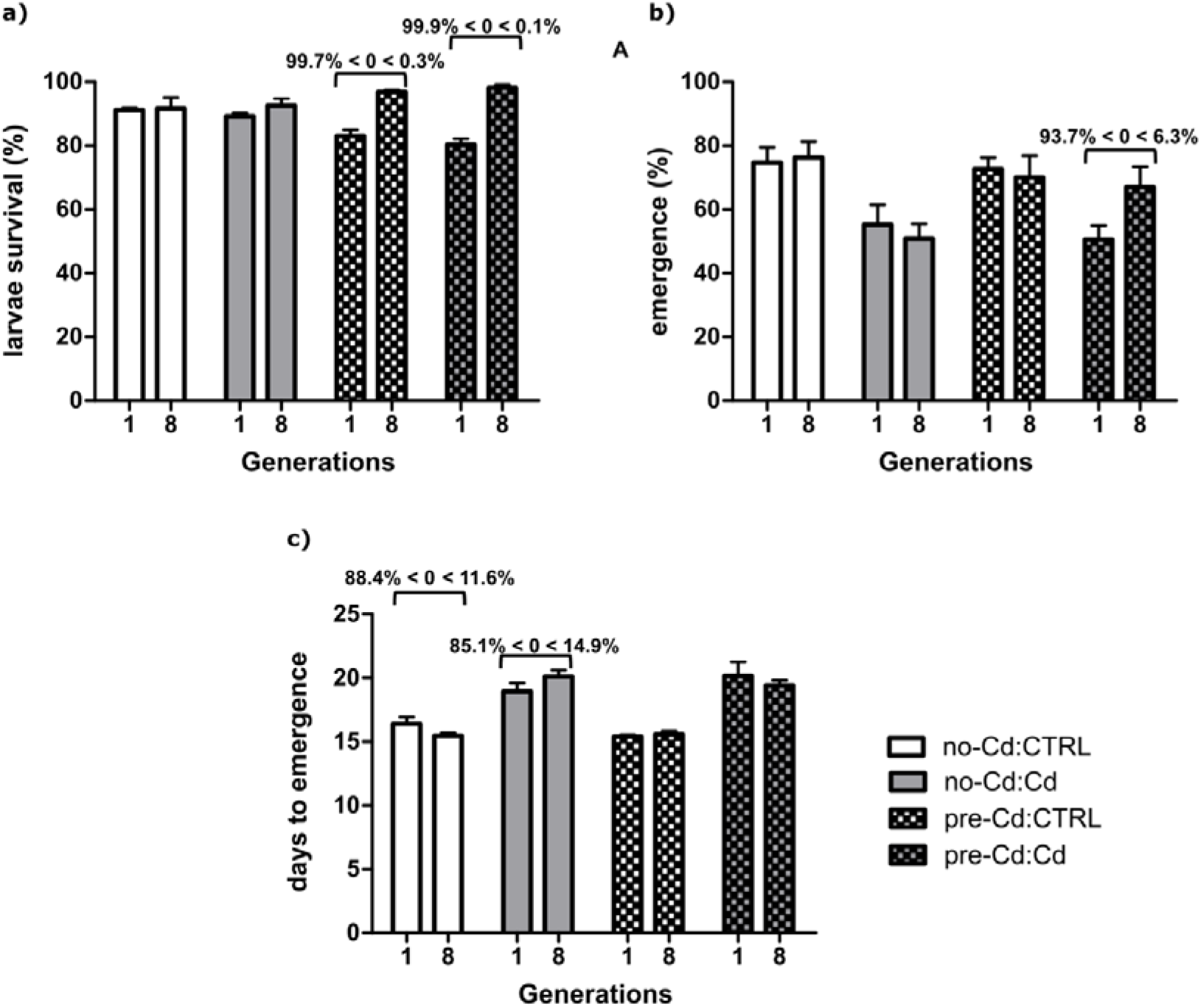
*C. riparius* life cycle measured endpoints comparing the first and eighth generations. a) Acute survival is estimated from the total of surving larvae after 24 hours. b) Chronic survival is estimated by the number of emerged adults. c) Median emergence time (EmT50) of *C. riparius*. For graphical representation pre-treatment is showed before (no-Cd or pre-Cd) and separated by a colon from the exposure condition (CTRL or Cd), that way a total of four groups (no-Cd:CTRL, no-Cd:Cd, pre-Cd:CTRL and pre-Cd:Cd) are showed. All bar graphs are expressed as mean values + SEM.

The recovery (REC) generation, which departed from the Cd pre-treated replicates on the sixth generation and were kept for two generations without further intended selective pressure regarding Cd, shows that the trend of improved survival in acute mortality endpoint was gained before the eighth generation and not lost during two generations without long-term exposure to Cd (Figure 3a). Both groups, exposed or not to Cd, showed that the difference of the means have more than 98% probability of being less than zero (difference of the means = −13 [−25, −2.9] and --15 [−22, −7.5]; probability of difference of the means = 98.2% < 0 < 1.8% and 99.6% < 0 < 0.4%, for CTRL and Cd exposed groups, respectively).

**Figure 3:**
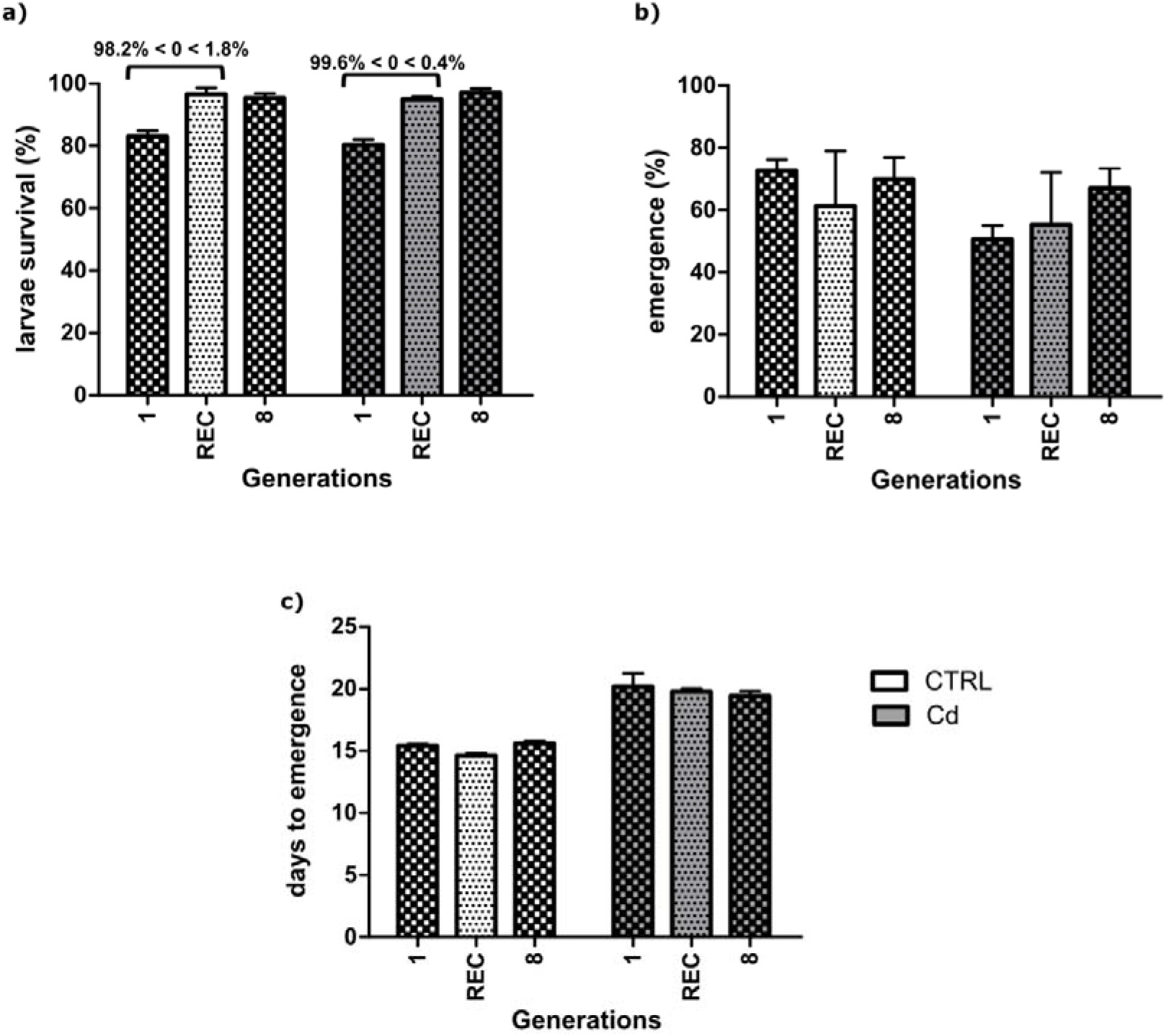
*C. riparius* measured endpoints comparing the recovery group (REC) to the first and eighth generation of the Cd pre-exposed groups (pre-Cd). Exposure to Cd to 0 (white) or 50 (grey) ug/L are showed as background colour. a) Acute survival is estimated from the total of surviving larvae after 24 hours. b) Chronic survival is estimated by the number of emerged adults. c) Median emergence time (EmT50) of *C. riparius*. All bar graphs are expressed as mean values ± SEM.

### 3.3. Time-series analysis of DNA Poolseq data

Effective population size from first to the most evolved generation did not differ statistically for control replicates (72±9.67) and for the Cd replicates (87.6±12). However, the control and Cd pre-treated replicates started at similar points, but after the third generation control replicates always showed higher population size than the Cd pre-treatment ones (Supplementary Information – S3, Figure 1). The mean AFC of the significant SNPs in the control replicates were 0.183±0.0464 and significantly lower (t = 4.3382, df = 47150, p-value = 1.44e-05) than in the Cd pre-treated replicates that showed 0.185±0.0455 (Supplementary Information – S3, Figure 3). Genetic diversity for the control replicates varied from 0.009975± 0.0001327 in the first generation to 0.01014± 0.0001118 in the most evolved generation. The Cd pre-treated replicates values fluctuated from 0,01004± 0,0002187, in the first generation, until 0,009918± 0,009918, in the last generation.

Regarding the time-series investigation of allele frequency changes it is certain that all the replicates in both groups experienced selective pressures, and therefore adapted to them, as many observed p-values were above the expected threshold found by the drift simulation (Figure 4a and 4b; Supplementary Information – S3, Table 1 and Figure 2). A total of 77,632 SNPs, 53,595 for the control replicates and 24,037 for the Cd pre-treated replicates, passed the filtering steps and were used for haplotype investigation.

**Figure 4:**
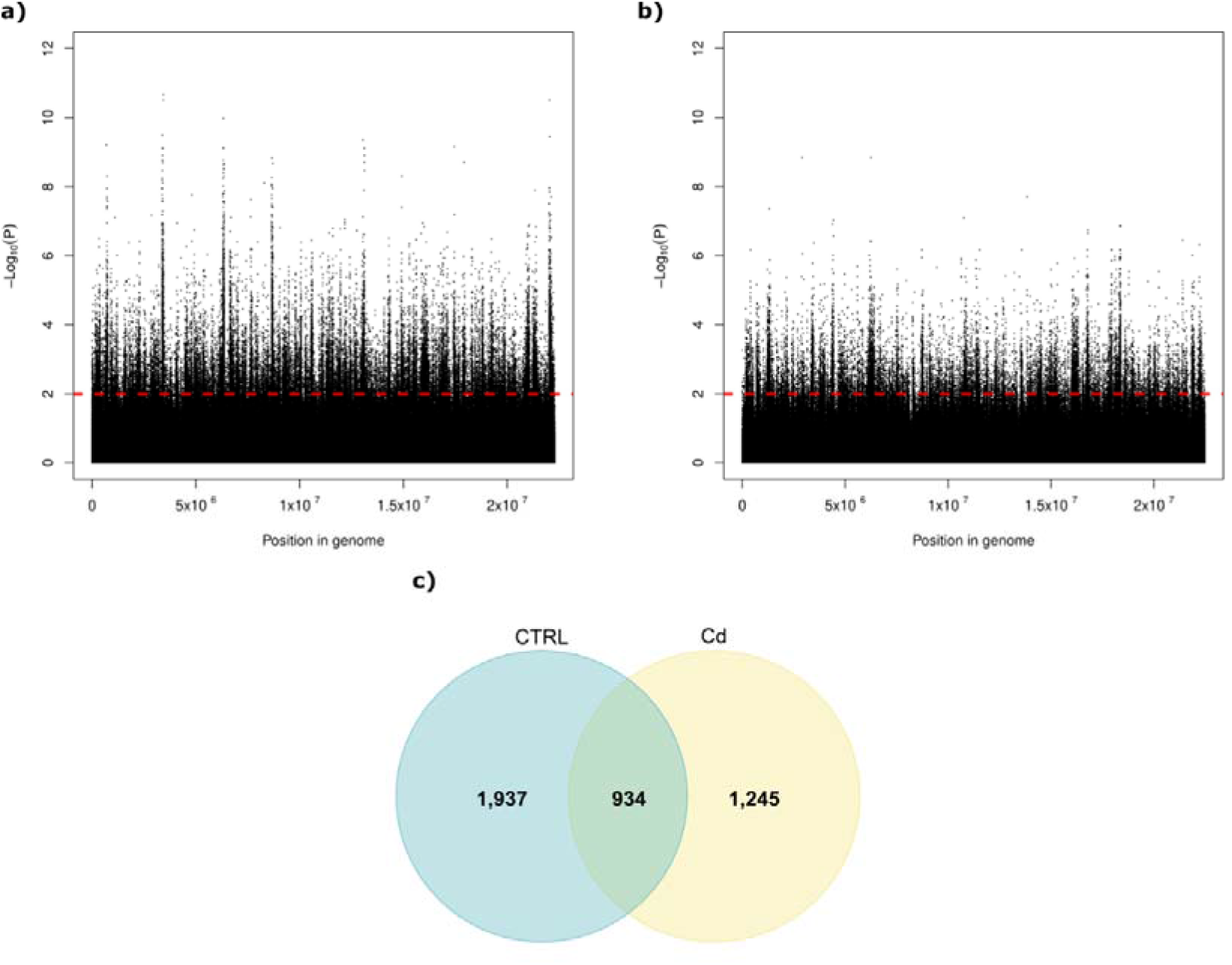
Manhattan plot of false discovery rate (BH) corrected -Log_10_ probability values from CMH test. a) for control replicates and b) for Cd pre-treated replicates. The dashed red horizontal line indicates the chosen significance threshold (p < 0.01) to consider SNPs as significant for the haplotype investigation. c) Venn diagram representing the results from haplotype investigation and genes of potential section target. The blue circle represents the genes identified in the control (CTRL) replicates and the yellow circle the ones involved in Cd pre-treatment (Cd).

From those filtered SNPs, it was possible to identify 6,668 haplotypes, 3,743 in the control replicates and 2,925 in the Cd pre-treated replicates, further, a total of 4,116 genes involved in both treatments were associated with the previously detected haplotypes. While 934 genes from the above-mentioned list were common to both groups, 1,937 genes appeared uniquely in the control group and 1,245 emerged in the Cd pre-treated group (Figure 4c). This resulted on 16 biological processes identified with a GO Term for the control and 9 for the Cd pre-treatment (Supplementary Information – S3, Table 2). Two biological processes (GO:0007186 and GO:0006355) representing G-protein coupled receptor signaling pathway and regulation of transcription are shared between the lists, also being the most representative ones in both lists. Other GO Terms overrepresented in both groups are connected to ion transmembrane transport (GO:0006813, GO:0055085, GO:0034220, GO:0006811, GO:0070588, GO:0008272).

**Table 2:** List of significantly enriched GO biological functions associated with the overlap between transcriptomic results and associated haplotypes in the Cd pre-treatment. GO terms were obtained using topGO with the wheight01 algorithm.

| GO.ID | Term | Annotated | Significant | Fisher |
| --- | --- | --- | --- | --- |
| GO:0060294 | cilium movement involved in cell motility | 1 | 1 | 0.0026 |
| GO:0007018 | microtubule-based movement | 54 | 3 | 0.0059 |
| GO:0048870 | cell motility | 4 | 2 | 0.0072 |

### 3.4. Gene expression analysis

An overview of the overall effect of experimental covariates (pre-treatment and treatment) on the groups are represented by a PCA (Figure 5a). Sample specific characteristics regarding expression patterns can be observed in a heatmap (Supplementary Information S4 - Figure 1). While the groups exposed to Cd clustered together independent from pre-exposure regime, when not challenged by Cd exposure they have different expressions patterns.

**Figure 5:**
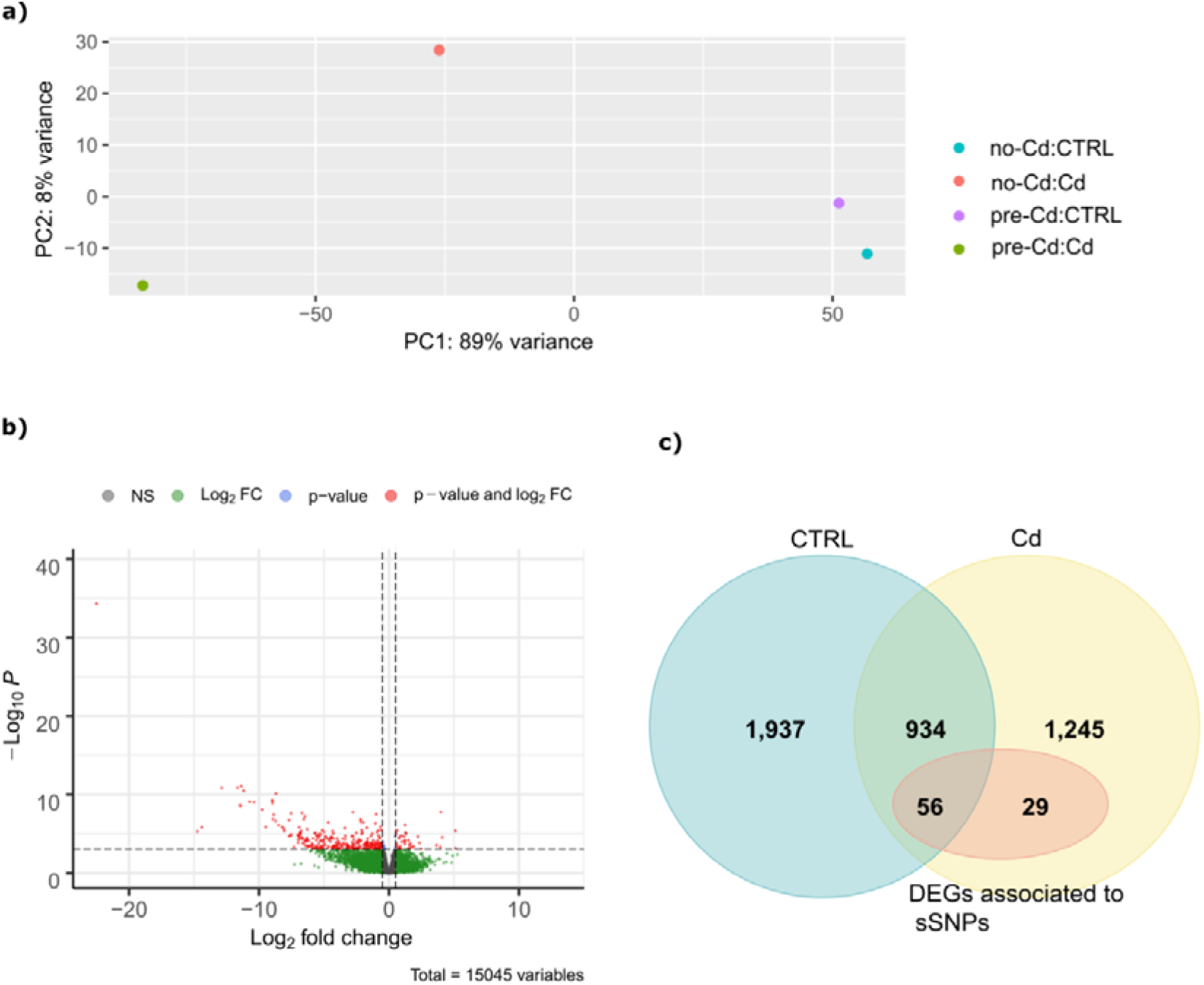
a) Gene expression PCA plot of the groups to visualize the overall effect of experimental covariates between the groups. Percentages on the x□ and y□axis indicate the percentage of variance explained by each principal component. b) Volcano plot for the DEGs found on the Cd pre-treatment. The threshold values are represented as dashed lines (p-value = 0.01, Fold change = 0.5). Genes above fold change and p-value threshold are colored in red. c) Venn diagram representing the intersection between the DNA Poolseq and transcriptome investigation results. The blue circle represents the genes identified in the control (CTRL) replicates and the yellow circle the ones involved in Cd pre-treatment (Cd). The red ellipsis shows the overlap between the DEGs in transcriptomic data and the significant SPNs (sSNPs) related to in Cd pre-treatment.

The pre-exposure to Cd, namely the long-term transgenerational exposure, revealed 389 DEGs. From those 223 DEGs were identified with a GO term (Figure 5b). Seven biological processes were overrepresented, revealing enrichment of several functions related to movement of organelles or other cellular components via polymerization or depolymerization of microtubules (GO:0007017, GO:0007018) (Supplementary Information S4 - Table 1). The integration of the transcriptomic data with the significant SNPs associated with Cd pre-treatment showed that from the 398 DEGs, 85 of those could be correlated with a gene associated to a significant SNP. Further, an overlap of 29 genes uniquely linked to the Cd pre-treatment were detected (Figure 5c). Which is higher and highly significant (p < 0.001) than the probability to observe an overlap by chance (1000 randomizations; min = 1, max = 17). The result of the GO Term enrichment points that microtubule movement and cell motility (GO:0007018, GO:0048870) were the two significant overrepresented terms of this intersection (Table 2).

The exposure to Cd during one life-cycle showed 2741 DEGs, from which sixteen significantly enriched biological process were identified. The most representative ones were linked to energy production, mainly ATP synthesis (GO:0015986, GO:0006099, GO:0006120, GO:0006122, GO:0022900), oxidative stress (GO:0055114) and visual and light perception (GO:0007601, GO:0007601) (Supplementary Information S4- Table 1). Further, although the LRT model revealed twenty-seven differentially expressed genes (DEGs) on the interaction between pre-treatment and exposure, none of those represented significantly enriched biological process.

## 4. DISCUSSION

To answer our main question if Cd tolerance acquisition is indeed heritable and based on genetic adaptation, we made use of a multigenerational approach in which the Cd concentration gradually increased over time during eight generations. We started with the current LOEC sediment concentration for Cd on *C. riparius* (Doria and Pfenninger, 2021), and finished with about 1.3 mg/kg of Cd, which represents a deleterious concentration for the species (Vogt et al., 2010), even though environmental concentrations are often higher. This particular scenario was designed to take into account the relevance of gradual directional environmental changes (Gorter et al., 2017), that can profoundly affect the course and outcome of evolution. Indeed, while consistent Cd tolerance can be observed on laboratory and field populations suddenly exposed to a constant high concentration of the metal (Chaumot et al., 2016; Pedrosa et al., 2017b; Postma and Davids, 1995; Ward and Robinson, 2005), Cd tolerance on populations exposed to sublethal concentrations is highly variable, occurring only in a small number of individuals and weakly heritable (Chaumot et al., 2016; Groenendijk et al., 2002; Marinković et al., 2012; Sheir et al., 2013).

Our results from the measured life-cycle endpoints endorse previous studies that decreased Cd sensitivity in *C. riparius* exposed to sublethal concentrations might be hard to detect (Doria and Pfenninger, 2021; Marinković et al., 2012; Vogt et al., 2010). This could be due to the limited duration of laboratory experiments in general; highly significant increases of fitness may take appreciably more generations to evolve. But, it is important to highlight that survival and developmental time are fitness factors that tend to possess low heritability compared to behavioural, morphological and physiological traits (Mousseau and Roff, 1987). Also, because those measurable life-cycle traits may have greater and variable nonadditive genetic sources of variance, differential response within fitness components to selection is known (Klemme and Hanski, 2009; Ohtsuki et al., 2020). In this regard, EmT50 had no alteration over the course of time (Doria and Pfenninger, 2021; Marinković et al., 2012, 2011; Postma and Davids, 1995) and continued to be equally higher in both groups exposed do Cd independent of pre-exposure scenario. However, the bayesian approach revealed convincing evidence that both acute and chronic survival improved under selection.

The acute mortality, assessed by newly hatched larval survival, uncovered the fact that larval survival grew over time independently from the exposure scenario in pre-treated replicates. This fitness component increased to the level of the control groups and remained stable even after two generations without Cd pre-treatment, which likely represents the maximum this population can achieve. Moreover, our findings are consistent with previous reports that *C. riparius* larvae are extremely resistant to Cd (Gillis and Wood, 2008).

Regarding the DNA time-series investigation, it is important to notice that the microevolution observation of allele frequency changes that occurred over time within the replicates, fundamentally occurred by selection since gene flow, genetic drift and elevated mutation rates (Doria et al., 2021) can be excluded. Hence, it is remarkable that although the final lists underlined allele frequency changes in genes unique to each treatment, those different selection targets still converged into two shared biological functions. Both overrepresented terms, namely the G-protein coupled receptor signaling pathway and the regulation of transcription, are connected and involved in sensing environmental cues and adjust numerous physiological processes like reproduction and feeding patterns (Li et al., 2014; Liu et al., 2021). They are also implicated in insect’s steroid hormone signaling that control growth and development (Liu et al., 2021; Wang et al., 2015). Thus, being involved in normal development of the chironomids.

This situation makes it is possible to affirm that even the control condition without exposure to any toxicant can also be considered a treatment to which the animals adapt continually over time. Highlighting that adaptation to laboratory conditions is always ongoing in multigenerational experiments of populations with genetic variation. But, although it was previously described for *C. riparius* (Pfenninger and Foucault, 2020b), it is still not considered as an influential factor in ecotoxicological bioassays studies, that focus more in effects of inbreeding in laboratory animals (Nowak et al., 2007).

Nonetheless, this adaptative evolution of the organisms to the laboratory conditions, could have masked the underlying Cd adaptation experienced by the population. Currently there is little consensus on the optimization of E&R studies to either maximize the power to detect selected variants or distinguish neutrally evolving loci from the ones selected to specific selective pressures (Otte and Schlötterer, 2021; Pfenninger and Foucault, 2020b). In our study we showed that integration of transcriptomic responses to time-series analysis of allele frequency changes is a good and reliable technique to pinpoint the genetic footprint exerted by exposure to low concentrations of Cd.

The overlap between DEGs and sSNPs unique to Cd pre-treatment showed that genes related to microtubules movement and cell motility had significant changes in their allele trajectories, which ultimately reflected on their expression patterns. This contradicts more recent data and empirically demonstrate that low environmental Cd concentrations can promote genetic adaptation (Doria and Pfenninger, 2021; Marinković et al., 2012; Vogt et al., 2010).

Microtubules organization, ultimately responsible for transepithelial transport events, intracellular trafficking and cellular excretion, are often altered in cells exposed to toxic metals and have been linked to metal tolerance in chironomids (Choong et al., 2014; Mattingly et al., 2001; Mireji et al., 2010; Wang et al., 2016). Therefore, this overrepresentation on the pre-treated group might reflect an attempt to have an efficient shedding of accumulated metals, already showed to happened in natural populations of chironomids exposed to metals (Dick Groenendijk et al., 1999). Interestingly, these findings can be implicated in our results from the life-cycle experiment, in which the chronic survival and EmT50 parameters indicated lower emergence rate and greater developmental time. Meaning that Cd has pronounced effects on metamorphosis success. This might be due to the great affinity of chironomids to bioaccumulate Cd during its larval stages and remove the metal during pupal stage with high rates of elimination on the exuviae (Dick Groenendijk et al., 1999; Timmermans and Walker, 1989). In the course of metamorphosis the imago develop from small groups of undifferentiated cells, already present in the larva but not functional, while most larval cells disintegrate (Timmermans and Walker, 1989). This disintegration allied to an efficient efflux of Cd can be an important metal elimination step. On the other hand, when this mechanism fails, it allows great concentrations of Cd to be bioavailable to the new cells forming the pupae and the adults. But, as the Cd effects are too strong for some organisms they may reach a tolerance threshold with mostly fatal consequences (Dallinger and Höckner, 2013), reflected by the lower emergence rates on the exposed groups irrespective of pre-treatment.

However, other metal responsive genes, like the metal-binding metallothioneins, that have been described to have a crucial role in reduced sensitivity to Cd in *C. riparius* and daphnids (Shaw et al., 2019; Toušová et al., 2016) have not been identified in our integrative analysis. Tolerant animals normally showed higher expression patterns of this genes and therefore, have a better intracellular regulation of metals (Mireji et al., 2010; Toušová et al., 2016). This might be due either to the genetic diversity and background available on the studied population or, most interesting, to our approach of temporal increasing concentrations of Cd in the replicates. Organisms exposed to low concentrations of a toxic metal may compensate the stress by changing normal physiological mechanisms such as digestive functions, absorption efficiencies, or pumping mechanisms at the cellular level (Dallinger and Höckner, 2013), which is in agreement with our findings.

Therefore, even if there was an effort to better eliminate Cd, it was not enough to avoid classical side effects of Cd exposure that primarily affects energy production pathways and promotes oxidative stress (Martín-Folgar and Martínez-Guitarte, 2019; Nair et al., 2013; Park et al., 2012; Pedrosa et al., 2017b). This situation occurs primarily because, to maintain physiological homeostasis, the stress imposed by Cd exposure generates the need to allocate more energy resources for defense against oxidative stress and other molecular damages (Pedrosa et al., 2017b). Interestingly, visual and light perception upon exposure to Cd were some overrepresented biological functions found on our results that are still hardly discussed in the literature. These pathways might be related either with the role of Cd on circadian disruption through oxidative stress and immune response (Jiménez-Ortega et al., 2011; Xiao et al., 2016), and therefore having an altered perception of the environment (Doria et al., 2018). As well as the differences on Cd damage extent between the dark and the light phase (Xiao et al., 2016).

Finally, our results suggest that Cd can promote genetic adaptation in exposed population. However, the adaptation footprint detected on the present study had only marginal effects on measured life-cycle endpoints. That raises the possibility that although *C. riparius* can adapt rapidly to environmental stress conditions (Foucault et al., 2018), metal sensitivity is a trait that slowly builds up throughout generations and might be marginally detectable if populations are not historically and heavily exposed to Cd (Groenendijk et al., 1999; Marinković et al., 2012; Pedrosa et al., 2017a). Importantly, we could successfully establish a multi-genomic approach to study and reliably detect adaptation when organisms are exposed to chemicals.

## 5. CONCLUSION

This is the first study to explore next generation sequencing within the context of adaptation to chemicals, more specifically, Cd. Using life-cycle endpoints, transcriptomic and population genetic analysis our results showed heritable genetic acquired resistance to Cd. Evidence of genetic adaptation was found only in a joint analysis of transcriptomic data and time-series analysis of E&R experiment. The main target of found adaptation relied on genes related to efflux of metals.

## REFERENCES

Alexa, A., Rahnenfuhrer, J., 2016. Gene set enrichment analysis with topGO. Bioconductor Improv.

Anders, S., Pyl, P.T., Huber, W., 2015. HTSeq-A Python framework to work with high-throughput sequencing data. Bioinformatics 31, 166–169. https://doi.org/10.1093/bioinformatics/btu638

Andrews, S., 2010. FastQC: A Quality Control Tool for High Throughput Sequence Data [WWW Document]. URL http://www.bioinformatics.babraham.ac.uk/projects/fastqc/

Bååth, R., 2014. Bayesian First Aid□: A Package that Implements Bayesian Alternatives to the Classical *. test Functions in R. Proc. UseR 2014 33, 2.

Barrett, R.D.H., Schluter, D., 2008. Adaptation from standing genetic variation. Trends Ecol. Evol. 23, 38–44. https://doi.org/10.1016/j.tree.2007.09.008

Beaudouin, R., Dias, V., Bonzom, J.M., Péry, A., 2012. Individual-based model of Chironomus riparius population dynamics over several generations to explore adaptation following exposure to uranium-spiked sediments. Ecotoxicology 21, 1225–1239. https://doi.org/10.1007/s10646-012-0877-4

Becker, J.M., Liess, M., 2015. Biotic interactions govern genetic adaptation to toxicants. Proc. R. Soc. B Biol. Sci. 282. https://doi.org/10.1098/rspb.2015.0071

Blickley, T.M., Matson, C.W., Vreeland, W.N., Rittschof, D., Di Giulio, R.T., McClellan-Green, P.D., 2014. Dietary CdSe/ZnS quantum dot exposure in estuarine fish: Bioavailability, oxidative stress responses, reproduction, and maternal transfer. Aquat. Toxicol. 148, 27–39. https://doi.org/10.1016/j.aquatox.2013.12.021

Blighe K, Rana S, L.M., 2021. EnhancedVolcano: Publication-ready volcano plots with enhanced colouring and labeling.

Bolger, A.M., Lohse, M., Usadel, B., 2014. Trimmomatic: A flexible trimmer for Illumina sequence data. Bioinformatics 30, 2114–2120. https://doi.org/10.1093/bioinformatics/btu170

Canzler, S., Schor, J., Busch, W., Schubert, K., Rolle-Kampczyk, U.E., Seitz, H., Kamp, H., von Bergen, M., Buesen, R., Hackermüller, J., 2020. Prospects and challenges of multi-omics data integration in toxicology. Arch. Toxicol. 94, 371–388. https://doi.org/10.1007/s00204-020-02656-y

Carrasco-Navarro, V., Muñiz-González, A.-B., Sorvari, J., Martínez-Guitarte, J.-L., 2021. Altered gene expression in Chironomus riparius (insecta) in response to tire rubber and polystyrene microplastics ⍰. Environ. Pollut. 285. https://doi.org/10.1016/j.envpol.2021.117462

Charlesworth, B., 2009. Effective population size and patterns of molecular evolution and variation. Nat. Rev. Genet. 10, 195–205. https://doi.org/10.1038/nrg2526

Chaumot, A., Gos, P., Garric, J., Geffard, O., 2009. Additive vs non-additive genetic components in lethal cadmium tolerance of Gammarus (Crustacea): Novel light on the assessment of the potential for adaptation to contamination. Aquat. Toxicol. 94, 294–299. https://doi.org/10.1016/j.aquatox.2009.07.015

Chaumot, A., Vigneron, A., Geffard, O., 2016. Mothers and not genes determine inherited differences in cadmium sensitivities within unexposed populations of the freshwater crustacean Gammarus fossarum. Evol. Appl. 355–366. https://doi.org/10.1111/eva.12327

Choong, G., Liu, Y., Templeton, D.M., 2014. Interplay of calcium and cadmium in mediating cadmium toxicity. Chem. Biol. Interact. 211, 54–65. https://doi.org/10.1016/j.cbi.2014.01.007

Dallinger, R., Höckner, M., 2013. Evolutionary concepts in ecotoxicology□: tracing the genetic background of differential cadmium sensitivities in invertebrate lineages. Ecotoxicology 767–778. https://doi.org/10.1007/s10646-013-1071-z

Dong, Q., Liu, Y., Liu, G., Guo, Y., Yang, Q., 2021. Enriched isotope tracing to reveal the fractionation and lability of legacy and newly introduced cadmium under different amendments. J. Hazard. Mater. 403, 123975. https://doi.org/10.1016/j.jhazmat.2020.123975

Doria, H.B., Ferreira, M.B., Rodrigues, S.D., Lo, S.M., Domingues, C.E., Nakao, L.S., de Campos, S.X., Ribeiro, C.A.D.O., Randi, M.A.F., 2018. Time does matter! Acute copper exposure abolishes rhythmicity of clock gene in Danio rerio. Ecotoxicol. Environ. Saf. 155. https://doi.org/10.1016/j.ecoenv.2018.02.068

Doria, H.B., Pfenninger, M., 2021. A multigenerational approach can detect early Cd pollution in Chironomus riparius. Chemosphere 262, 127815.https://doi.org/10.1016/j.chemosphere.2020.127815

Doria, H.B., Waldvogel, A.M., Pfenninger, M., 2021. Measuring mutagenicity in ecotoxicology: A case study of Cd exposure in Chironomus riparius. Environ. Pollut. 272, 116004. https://doi.org/10.1016/j.envpol.2020.116004

Ebner, J.N., 2021. Trends in the Application of “Omics” to Ecotoxicology and Stress Ecology. Genes (Basel). 12. https://doi.org/10.1007/978-94-007-2072-5

Foucault, Q., Wieser, A., Waldvogel, A.M., Feldmeyer, B., Pfenninger, M., 2018. Rapid adaptation to high temperatures in Chironomus riparius. Ecol. Evol. 8, 12780–12789. https://doi.org/10.1002/ece3.4706

Foucault, Q., Wieser, A., Waldvogel, A.M., Pfenninger, M., 2019. Establishing laboratory cultures and performing ecological and evolutionary experiments with the emerging model species Chironomus riparius. J. Appl. Entomol. 1–9. https://doi.org/10.1111/jen.12606

Gillis, P.L., Wood, C.M., 2008. The effect of extreme waterborne cadmium exposure on the internal concentrations of cadmium, calcium, and sodium in Chironomus riparius larvae. Ecotoxicol. Environ. Saf. 71, 56–64. https://doi.org/10.1016/j.ecoenv.2007.08.003

Gorter, F.A., Derks, M.F.L., Van Den Heuvel, J., Aarts, M.G.M., Zwaan, B.J., De Ridder, D., De Visser, J.A.G.M., 2017. Genomics of Adaptation Depends on the Rate of Environmental Change in Experimental Yeast Populations. Mol. Biol. Evol. 34, 2613–2626. https://doi.org/10.1093/molbev/msx185

Groenendijk, Dick, Kraak, M.H.S., Admiraal, W., 1999. Efficient shedding of accumulated metals during metamorphosis in metal-adapted populations of the midge Chironomus riparius. Environ. Toxicol. Chem. 18, 1225–1231. https://doi.org/10.1002/etc.5620180622

Groenendijk, D., Lücker, S.M.G., Plans, M., Kraak, M.H.S., Admiraal, W., 2002. Dynamics of metal adaptation in riverine chironomids. Environ. Pollut. 117, 101–109. https://doi.org/10.1016/S0269-7491(01)00154-3

Groenendijk, D, Opzeeland, B. Van, Pires, L.M.D., Postma, J.F., 1999. Fluctuating Life-History Parameters Indicating Temporal Variability in Metal Adaptation in Riverine Chironomids. Arch. Environ. Contam. Toxicol. 181, 175–181.

Horváth, G., Móra, A., Bernáth, B., Kriska, G., 2011. Polarotaxis in non-biting midges: Female chironomids are attracted to horizontally polarized light. Physiol. Behav. 104, 1010–1015. https://doi.org/10.1016/j.physbeh.2011.06.022

Im, J., Chatterjee, N., Choi, J., 2019. Genetic, epigenetic, and developmental toxicity of Chironomus riparius raised in metal-contaminated field sediments: A multi-generational study with arsenic as a second challenge. Sci. Total Environ. 672, 789–797. https://doi.org/10.1016/j.scitotenv.2019.04.013

Jiménez-Ortega, V., Cano-Barquilla, P., Scacchi, P.A., Cardinali, D.P., Esquifino, A.I., 2011. Cadmium-induced disruption in 24-h expression of clock and redox enzyme genes in rat medial basal hypothalamus: Prevention by melatonin. Front. Neurol. MAR, 1–9. https://doi.org/10.3389/fneur.2011.00013

Jorde, P.E., Ryman, N., 2007. Unbiased estimator for genetic drift and effective population size. Genetics 177, 927–935. https://doi.org/10.1534/genetics.107.075481

Kim, D., Paggi, J.M., Park, C., Bennett, C., Salzberg, S.L., 2019. Graph-based genome alignment and genotyping with HISAT2 and HISAT-genotype. Nat. Biotechnol. 37, 907–915. https://doi.org/10.1038/s41587-019-0201-4

Klemme, I., Hanski, I., 2009. Heritability of and strong single gene (Pgi) effects on life-history traits in the Glanville fritillary butterfly. J. Evol. Biol. 22, 1944–1953. https://doi.org/10.1111/j.1420-9101.2009.01807.x

Klerks, P.L., Weis, J.S., 1987. Genetic adaptation to heavy metals in aquatic organisms: A review. Environ. Pollut. 45, 173–205. https://doi.org/10.1016/0269-7491(87)90057-1

Klerks, P.L., Xie, L., Levinton, J.S., 2011. Quantitative genetics approaches to study evolutionary processes in ecotoxicology □; a perspective from research on the evolution of resistance 513–523. https://doi.org/10.1007/s10646-011-0640-2

Kofler, R., Orozco-terwengel, P., Maio, N. De, Pandey, R.V., Nolte, V., Kosiol, C., Schlo, C., 2011a. PoPoolation□: A Toolbox for Population Genetic Analysis of Next Generation Sequencing Data from Pooled Individuals 6. https://doi.org/10.1371/journal.pone.0015925

Kofler, R., Pandey, R.V., Schlötterer, C., Populationsgenetik, I., Vienna, V., Wien, A.-, 2011b. PoPoolation2□: identifying differentiation between populations using sequencing of pooled DNA samples (Pool-Seq) 27, 3435–3436. https://doi.org/10.1093/bioinformatics/btr589

Kofler, R., Schlötterer, C., 2014. A guide for the design of evolve and resequencing studies. Mol. Biol. Evol. 31, 474–483. https://doi.org/10.1093/molbev/mst221

Kolde, R., 2019. pheatmap: Pretty Heatmaps.

Kruschke, J.K., 2013. Bayesian estimation supersedes the t test. J. Exp. Psychol. Gen. 142, 573–603. https://doi.org/10.1037/a0029146

Li, H., Durbin, R., 2009. Fast and accurate short read alignment with Burrows-Wheeler transform. Bioinformatics 25, 1754–1760. https://doi.org/10.1093/bioinformatics/btp324

Li, H., Handsaker, B., Wysoker, A., Fennell, T., Ruan, J., Homer, N., Marth, G., Abecasis, G., Durbin, R., Data, G.P., Sam, T., 2009. The Sequence Alignment / Map format and SAMtools 25, 2078–2079. https://doi.org/10.1093/bioinformatics/btp352

Li, T., Liu, L., Zhang, L., Liu, N., 2014. Role of G-protein-coupled receptor-related genes in insecticide resistance of the mosquito, Culex quinquefasciatus. Sci. Rep. 4, 1–9. https://doi.org/10.1038/srep06474

Liu, N., Wang, Y., Li, T., Feng, X., 2021. G-protein coupled receptors (Gpcrs): Signaling pathways, characterization, and functions in insect physiology and toxicology. Int. J. Mol. Sci. 22, 14–18. https://doi.org/10.3390/ijms22105260

Long, A., Liti, G., Luptak, A., Tenaillon, O., 2015. Elucidating the molecular architecture of adaptation via evolve and resequence experiments. Nat. Rev. Genet. 16, 567–582. https://doi.org/10.1038/nrg3937

Love, M.I., Huber, W., Anders, S., 2014. Moderated estimation of fold change and dispersion for RNA-seq data with DESeq2. Genome Biol. 15, 1–21. https://doi.org/10.1186/s13059-014-0550-8

Marini, F., Binder, H., 2019. PcaExplorer: An R/Bioconductor package for interacting with RNA-seq principal components. BMC Bioinformatics 20, 1–8. https://doi.org/10.1186/s12859-019-2879-1

Marinković. M., De Bruijn, K., Asselman, M., Bogaert, M., Jonker, M.J., Kraak, M.H.S., Admiraal, W., 2012. Response of the nonbiting midge Chironomus riparius to multigeneration toxicant exposure. Environ. Sci. Technol. 46, 12105–12111. https://doi.org/10.1021/es300421r

Marinković, M., Verweij, R.A., Nummerdor, G.A., Jonker, M.J., Kraak, M.H.S., Admiraal, W., 2011. Life cycle responses of the midge chironomus riparius to compounds with different modes of action. Environ. Sci. Technol. 45, 1645–1651.https://doi.org/10.1021/es102904y

Martín-Folgar, R., Martínez-Guitarte, J.L., 2019. Effects of single and mixture exposure of cadmium and copper in apoptosis and immune related genes at transcriptional level on the midge Chironomus riparius Meigen (Diptera, Chironomidae). Sci. Total Environ. 677, 590–598. https://doi.org/10.1016/j.scitotenv.2019.04.364

Mattingly, K.S., Beaty, B.J., Mackie, R.S., McGaw, M., Carlson, J.O., Rayms-Keller, A., 2001. Molecular cloning and characterization of a metal responsive Chironomus tentans alphatubulin cDNA. Aquat. Toxicol. 54, 249–260. https://doi.org/10.1016/S0166-445X(00)00181-8

Mireji, P.O., Keating, J., Hassanali, A., Impoinvil, D.E., Mbogo, C.M., Muturi, M.N., Nyambaka, H., Kenya, E.U., Githure, J.I., Beier, J.C., 2010. Expression of metallothionein and α-tubulin in heavy metal-tolerant Anopheles gambiae sensu stricto (Diptera: Culicidae). Ecotoxicol. Environ. Saf. 73, 46–50. https://doi.org/10.1016/j.ecoenv.2009.08.004

Mousseau, T.A., Roff, D.A., 1987. Natural selection and the heritability of fitness components. Heredity (Edinb). 59, 181–197.

Nair, P.M.G., Park, S.Y., Choi, J., 2013. Characterization and expression of cytochrome p450 cDNA (CYP9AT2) in Chironomus riparius fourth instar larvae exposed to multiple xenobiotics. Environ. Toxicol. Pharmacol. 36, 1133–1140. https://doi.org/10.1016/j.etap.2013.08.011

Nowak, C., Jost, D., Vogt, C., Oetken, M., Schwenk, K., Oehlmann, J., 2007. Consequences of inbreeding and reduced genetic variation on tolerance to cadmium stress in the midge Chironomus riparius. Aquat. Toxicol. 85, 278–284. https://doi.org/10.1016/j.aquatox.2007.04.015

Nowak, C., Vogt, C., Pfenninger, M., Schwenk, K., Oehlmann, J., Streit, B., Oetken, M., 2009. Rapid genetic erosion in pollutant-exposed experimental chironomid populations. Environ. Pollut. 157, 881–886. https://doi.org/10.1016/j.envpol.2008.11.005

OECD, 2011. Test No. 235: Chironomus sp., Acute Immobilisation Test 1–17.

OECD, 2010. Test No. 233: Sediment-Water Chironomid Life-Cycle Toxicity Test Using Spiked Water or Spiked Sediment. https://doi.org/10.1787/9789264090910-en

Ohtsuki, H., Rueffler, C., Wakano, J.Y., Parvinen, K., Lehmann, L., 2020. The components of directional and disruptive selection in heterogeneous group-structured populations. J. Theor. Biol. 507, 110449. https://doi.org/10.1016/j.jtbi.2020.110449

Otte, K.A., Schlötterer, C., 2021. Detecting selected haplotype blocks in evolve and resequence experiments. Mol. Cell 93–109. https://doi.org/10.1111/1755-0998.13244

Park, S.Y., Nair, P.M.G., Choi, J., 2012. Characterization and expression of superoxide dismutase genes in Chironomus riparius (Diptera, Chironomidae) larvae as a potential biomarker of ecotoxicity. Comp. Biochem. Physiol. - C Toxicol. Pharmacol. 156, 187–194. https://doi.org/10.1016/j.cbpc.2012.06.003

Pauget, B., Gimbert, F., Coeurdassier, M., Crini, N., Pérès, G., Faure, O., Douay, F., Richard, A., Grand, C., Vaufleury, A. De, 2013. Assessing the in situ bioavailability of trace elements to snails using accumulation kinetics. Ecol. Indic. 34, 126–135. https://doi.org/10.1016/j.ecolind.2013.04.018

Pedrosa, J., Cocchiararo, B., Bordalo, M.D., Rodrigues, A.C., Soares, A.M.V.M., Barata, C., Nowak, C., Pestana, J., 2017a. The role of genetic diversity and past-history selection pressures in the susceptibility of Chironomus riparius populations to environmental stress. Sci. Total Environ. 576, 807–816. https://doi.org/10.1016/j.scitotenv.2016.10.100

Pedrosa, J., Gravato, C., Campos, D., Cardoso, P., Figueira, E., Nowak, C., Soares, A.M.V.M., Barata, C., Pestana, J.L.T., 2017b. Investigating heritability of cadmium tolerance in Chironomus riparius natural populations: A physiological approach. Chemosphere 170, 83–94. https://doi.org/10.1016/j.chemosphere.2016.12.008

Pfenninger, M., Foucault, Q., 2020a. Quantifying the selection regime in a natural Chironomus riparius population. bioRxiv 2020.06.16.154054. https://doi.org/10.1101/2020.06.16.154054

Pfenninger, M., Foucault, Q., 2020b. Genomic processes underlying rapid adaptation of a natural Chironomus riparius population to unintendedly applied experimental selection pressures. Mol. Ecol. 29, 536–548. https://doi.org/10.1111/mec.15347

Pfenninger, M., Reuss, F., Kiebler, A., Schönnenbeck, P., Caliendo, C., Gerber, S., Cocchiararo, B., Reuter, S., Blüthgen, N., Mody, K., Mishra, B., Bálint, M., Thines, M., Feldmeyer, B., 2021. Genomic basis for drought resistance in european beech forests threatened by climate change. Elife 10, 1–17. https://doi.org/10.7554/eLife.65532

Postma, J.F., Davids, C., 1995. Tolerance induction and life cycle changes in cadmium-exposed Chironomus riparius (Diptera) during consecutive generations. Ecotoxicol. Environ. Saf. https://doi.org/10.1006/eesa.1995.1024

Postma, J.F., Kleunen, A. Van, Admiraal, W., 1995. Alterations in Life-History Traits of Chironomus riparius (Diptera) Obtained from Metal Contaminated Rivers. Arch. Environ. Contam. Toxicol. 29, 469–475.

Postma, Jaap F, Kyed, M., Admiraal, W., 1995. Site specific differentiation in metal tolerance in the midge Chironomus riparius (Diptera, Chironomidae). Hydrobiologia 315, 159–165.

Rodríguez-Romero, A., Viguri, J.R., Calosi, P., 2021. Acquiring an evolutionary perspective in marine ecotoxicology to tackle emerging concerns in a rapidly changing ocean. Sci. Total Environ. 764, 142816. https://doi.org/10.1016/j.scitotenv.2020.142816

Schmidt, H., Hellmann, S.L., Waldvogel, A.M., Feldmeyer, B., Hankeln, T., Pfenninger, M., 2020. A high-quality genome assembly from short and long reads for the non-biting midge chironomus riparius (Diptera). Genes Genomes Genet. 10, 1151–1157. https://doi.org/10.3390/W11091751

Shaw, J.R., Colbourne, J.K., Glaholt, S.P., Turner, E., Folt, C.L., Chen, C.Y., 2019. Dynamics of Camium Acclimation in Daphnia pulex: Linking Fitness Costs, Cross-Tolerance, an Hyper-Inuction of Metallothionein. Environ. Sci. Technol. 53, 14670–14678. https://doi.org/10.1021/acs.est.9b05006

Sheir, S.K., Handy, R.D., Henry, T.B., 2013. Effect of pollution history on immunological responses and organ histology in the marine mussel mytilus edulis exposed to cadmium. Arch. Environ. Contam. Toxicol. 64, 701–716. https://doi.org/10.1007/s00244-012-9868-y

Straub, L., Strobl, V., Neumann, P., 2020. The need for an evolutionary approach to ecotoxicology. Nat. Ecol. Evol. 4, 895. https://doi.org/10.1038/s41559-020-1194-6

Taus, T., Futschik, A., Schlötterer, C., 2017. Quantifying Selection with Pool-Seq Time Series Data. Mol. Biol. Evol. 34, 3023–3034. https://doi.org/10.1093/molbev/msx225

Timmermans, K.R., Walker, P.A., 1989. The fate of trace metals during the metamorphosis of chironomids (diptera, chironomidae). Environ. Pollut. 62, 73–85. https://doi.org/10.1016/0269-7491(89)90097-3

Toušová, Z., Kuta, J., Hynek, D., Adam, V., Kizek, R., Bláha, L., Hilscherová, K., 2016. Metallothionein modulation in relation to cadmium bioaccumulation and age-dependent sensitivity of Chironomus riparius larvae. Environ. Sci. Pollut. Res. 23, 10504–10513. https://doi.org/10.1007/s11356-016-6362-5

Vogt, C., He, M., Nowak, C., Diogo, J.B., Oehlmann, J., Oetken, M., 2010. Effects of cadmium on life-cycle parameters in a multi-generation study with Chironomus riparius following a pre-exposure of populations to two different tributyltin concentrations for several generations. Ecotoxicology 19, 1174–1182. https://doi.org/10.1007/s10646-010-0501-4

Wang, D., Zhao, W.L., Cai, M.J., Wang, J.X., Zhao, X.F., 2015. G-protein-coupled receptor controls steroid hormone signaling in cell membrane. Sci. Rep. 5, 1–11. https://doi.org/10.1038/srep08675

Wang, S., Wu, W., Liu, F., 2019. Assessment of the human health risks of heavy metals in nine typical areas. Environ. Sci. Pollut. Res. 26, 12311–12323. https://doi.org/10.1007/s11356-018-04076-z

Wang, T., Wang, Q., Song, R., Zhang, Y., Yang, J., Wang, Y., Yuan, Y., Bian, J., Liu, X., Gu, J., Zhu, J., Liu, Z., 2016. Cadmium induced inhibition of autophagy is associated with microtubule disruption and mitochondrial dysfunction in primary rat cerebral cortical neurons. Neurotoxicol. Teratol. 53, 11–18. https://doi.org/10.1016/j.ntt.2015.11.007

Ward, T.J., Robinson, W.E., 2005. Evolution of Cadmium resistance in Daphnia magna. Environ. Toxicol. Chem. 24, 2341–2349.

Xiao, B., Chen, T.M., Zhong, Y., 2016. Possible molecular mechanism underlying cadmium-induced circadian rhythms disruption in zebrafish. Biochem. Biophys. Res. Commun. 481, 201–205. https://doi.org/10.1016/j.bbrc.2016.10.081

Zhang, C., Jansen, M., De Meester, L., Stoks, R., 2019. Rapid evolution in response to warming does not affect the toxicity of a pollutant: Insights from experimental evolution in heated mesocosms. Evol. Appl. 12, 977–988. https://doi.org/10.1111/eva.12772

