## Supplementary Information for "Detecting Cd adaptation footprint in C. riparius with a multi-genomic approach"

**S1 – Feeding Protocol**

| day mg/larvae mg in 50ml for 30 larvae | | |
| --- | --- | --- |
| 1 | 0.10 | 150 |
| 2 | 0.10 | 150 |
| 3 | 0.17 | 250 |
| 4 | 0.17 | 250 |
| 5 | 0.23 | 350 |
| 6 | 0.23 | 350 |
| 7 | 0.30 | 450 |
| 8 | 0.30 | 450 |
| 9 | 0.37 | 550 |
| 10 | 0.37 | 550 |
| 11 | 0.43 | 650 |
| 12 | 0.43 | 650 |
| 13 | 0.50 | 750 |
| 14 | 0.50 | 750 |
| 15 | 0.50 | 750 |
| 16 | 0.50 | 750 |
| 17 | 0.50 | 750 |
| 18 | 0.50 | 750 |
| 19 | 0.50 | 750 |
| 20 | 0.50 | 750 |
| 21 | 0.50 | 750 |
| 22 | 0.50 | 750 |
| 23 | 0.50 | 750 |
| 24 | 0.50 | 750 |
| 25 | 0.50 | 750 |
| 26 | 0.50 | 750 |
| 27 | 0.50 | 750 |
| 28 | 0.50 | 750 |

The larvae of each generation were fed according to the above protocol. Each morning 1 ml from a solution of deionized water and finely grounded fish food was delivered in the experimental bowls.

**S2 – t-test, ANOVA and p-values from Tuckey’s post hoc test for acute and chronic survival endpoints and the EmT50**

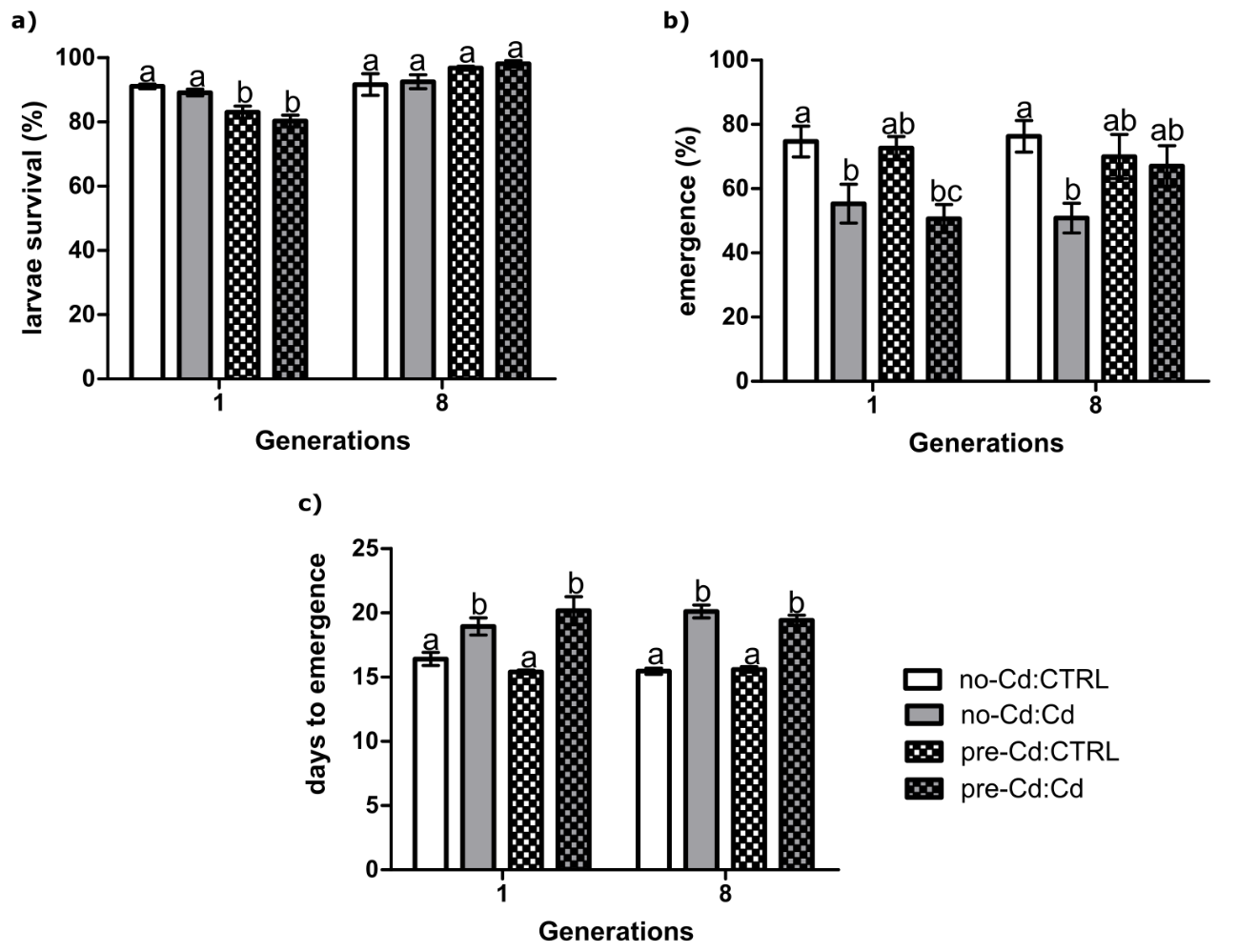

Figure 1: *C. riparius* life cycle measured endpoints. a) Acute survival is estimated from the total of surving larvae after 24 hours. b) Chronic survival is estimated by the number of emerged adults. c) Median emergence time (EmT50) of *C.* *riparius.* For graphical representation pre-treatment is showed before (no-Cd or pre-Cd) and separated by a colon from the exposure condition (CTRL or Cd), that way a total of four groups (no-Cd:CTRL, no-Cd:Cd, pre-Cd:CTRL and pre-Cd:Cd) are showed. All bar graphs are expressed as mean values ± SEM. Different letters indicate significant differences between groups of generations (p <0.05).

Table 1: ANOVA associated degrees of freedom, F values and p-values for acute and chronic mortality and EmT50 at the three analyzed generations.

| **generation**  **/endpoint** | | **df** | **F** | **p-value** |
| --- | --- | --- | --- | --- |
| **1** | **acute mortality** | 3 | 12.23 | 2.06e-04*** |
|  | **chronic mortality** | 3 | 6.439 | 0.0045** |
|  | **EmT50** | 3 | 10.07 | 5.73e-04*** |
| **8** | **acute mortality** | 3 | 2.251 | 0.101 |
|  | **chronic mortality** | 3 | 2.978 | 0.0455* |
|  | **EmT50** | 3 | 46.85 | 1.63e-12*** |

Asterisks indicate significant differences (***p<0.001; **p<0.01; *p<0.05).

Table 2: p-values of the Tuckey’s post hoc test for each comparison between the groups for acute and chronic mortality and EmT50 at the three analyzed generations.

| **generation/ groups** | | **acute mortality** | **chronic mortality** | **EmT50** |
| --- | --- | --- | --- | --- |
| **1** | **pre-Cd:CTRL/pre-Cd:Cd** | 0.5455 | 0.0229* | 9.1e-04*** |
|  | **no-Cd:Cd/pre-Cd:Cd** | 0.0026** | 0.8988 | 0.6078 |
|  | **no-Cd:CTRL/pre-Cd:Cd** | 4e-04*** | 0.0126* | 0.0072** |
|  | **no-Cd:Cd/pre-Cd:CTRL** | 0.04026* | 0.0866 | 0.0114* |
|  | **no-Cd:CTRL/pre-Cd:CTRL** | 0.0061** | 0.9905 | 0.7381 |
|  | **no-Cd:CTRL/no-Cd:Cd** | 0.7809 | 0.0496* | 0.0846 |
| **8** | **pre-Cd:CTRL/pre-Cd:Cd** | 0.9656 | 0.9825 | <2e-16*** |
|  | **no-Cd:Cd/pre-Cd:Cd** | 0.2582 | 0.2581 | 0.5467 |
|  | **no-Cd:CTRL/pre-Cd:Cd** | 0.1503 | 0.6841 | <2e-16*** |
|  | **no-Cd:Cd/pre-Cd:CTRL** | 0.5033 | 0.1381 | <2e-16*** |
|  | **no-Cd:CTRL/pre-Cd:CTRL** | 0.3335 | 0.8743 | 0.9924 |
|  | **no-Cd:CTRL/no-Cd:Cd** | 0.9899 | 0.0331* | <2e-16*** |

Asterisks indicate significant difference between the compared groups (***p<0.001; **p<0.01; *p<0.05).

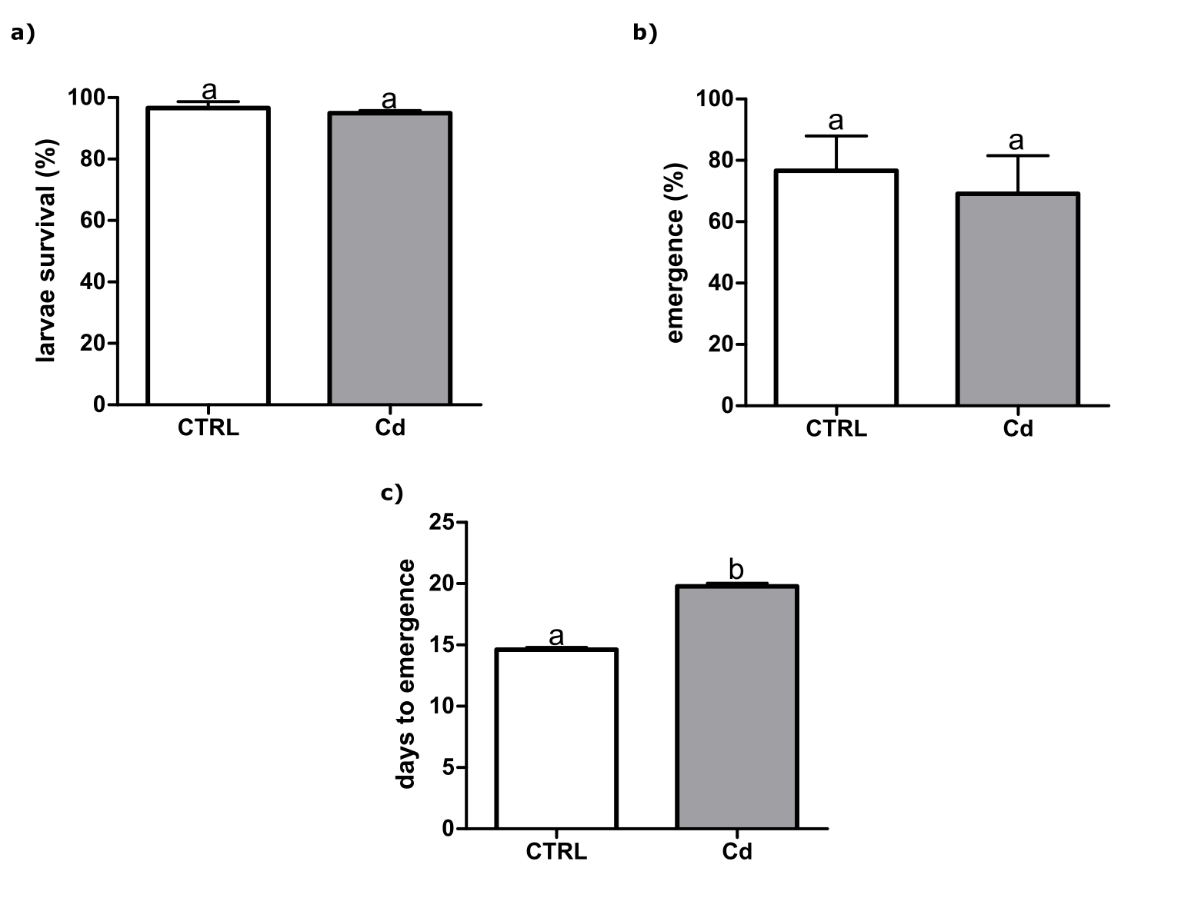

Figure 2: *C. riparius* measured endpoints of the recovery group (REC) exposed to 0 or 50 ug/L of Cd. a) Acute survival is estimated from the total of surviving larvae after 24 hours. b) Chronic survival is estimated by the number of emerged adults. c) Median emergence time (EmT50) of *C.* *riparius.* All bar graphs are expressed as mean values ± SEM. Different letters indicate significant differences between groups of generations (p <0.05).

Table 3: T-test t values, associated degrees of freedom, and p-values for the comparison between non-exposed (CTRL) and exposed (Cd) groups in the recovery (REC) set-up at the first and second generations

| **endpoint** | **t** | **df** | | **p-value** | |
| --- | --- | --- | --- | --- | --- |
| **acute mortality**  **chronic mortality**  **EmT50** | -0.7267 | | 8 | | 0.4881 |
|  | -0.24604 | | 8 | | 0.8118 |
|  | 16.372 | | 8 | | 3.31e-06*** |

Asterisks indicate significant difference between the comparisons (***p<0.001; *p<0.05).

**S3 – Time-series analysis of DNA Poolseq data**

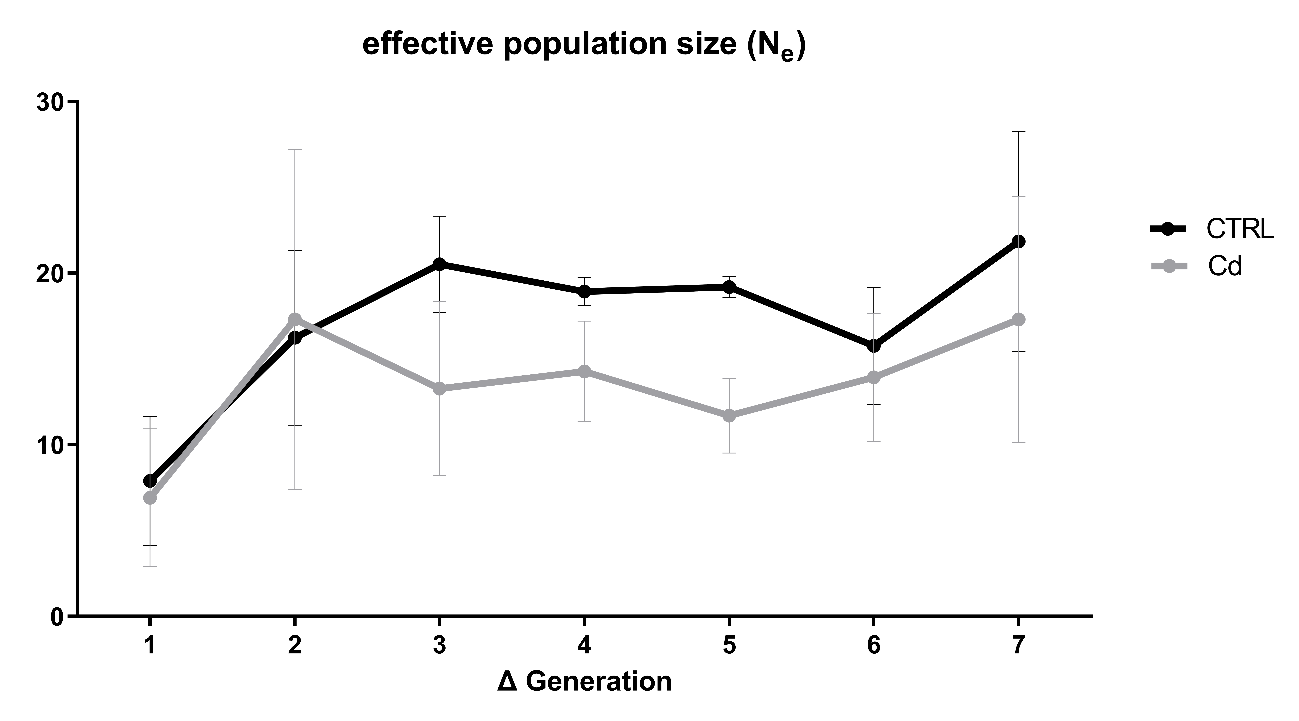

Figure 1: Effective population size measured by Watterson’s theta at every generational turn. Black line represents the control replicates (CTRL) and grey line represents the Cd pre-treated replicates (Cd).

Table 1: Corrected and transformed p-values for observed changes compared to the drift expectation on the three replicates of each group control (CTRL) and pre-treated with Cd (Cd).

| **group/**  **replicate** | **CTRL** | | **Cd** | |
| --- | --- | --- | --- | --- |
|  | **observed changes** | **expected drift** | **observed changes** | **expected drift** |
| **A** | 3.305072 | 2.983003 | 3.445553 | 2.740926 |
| **B** | 3.964317 | 3.638796 | 2.859201 | 1.983248 |
| **C** | 3.070031 | 2.427724 | 2.329976 | 1.7309987 |

A quantile of 99.90% was set, representing the threshold for highly significant SNPs according to chi-square test.

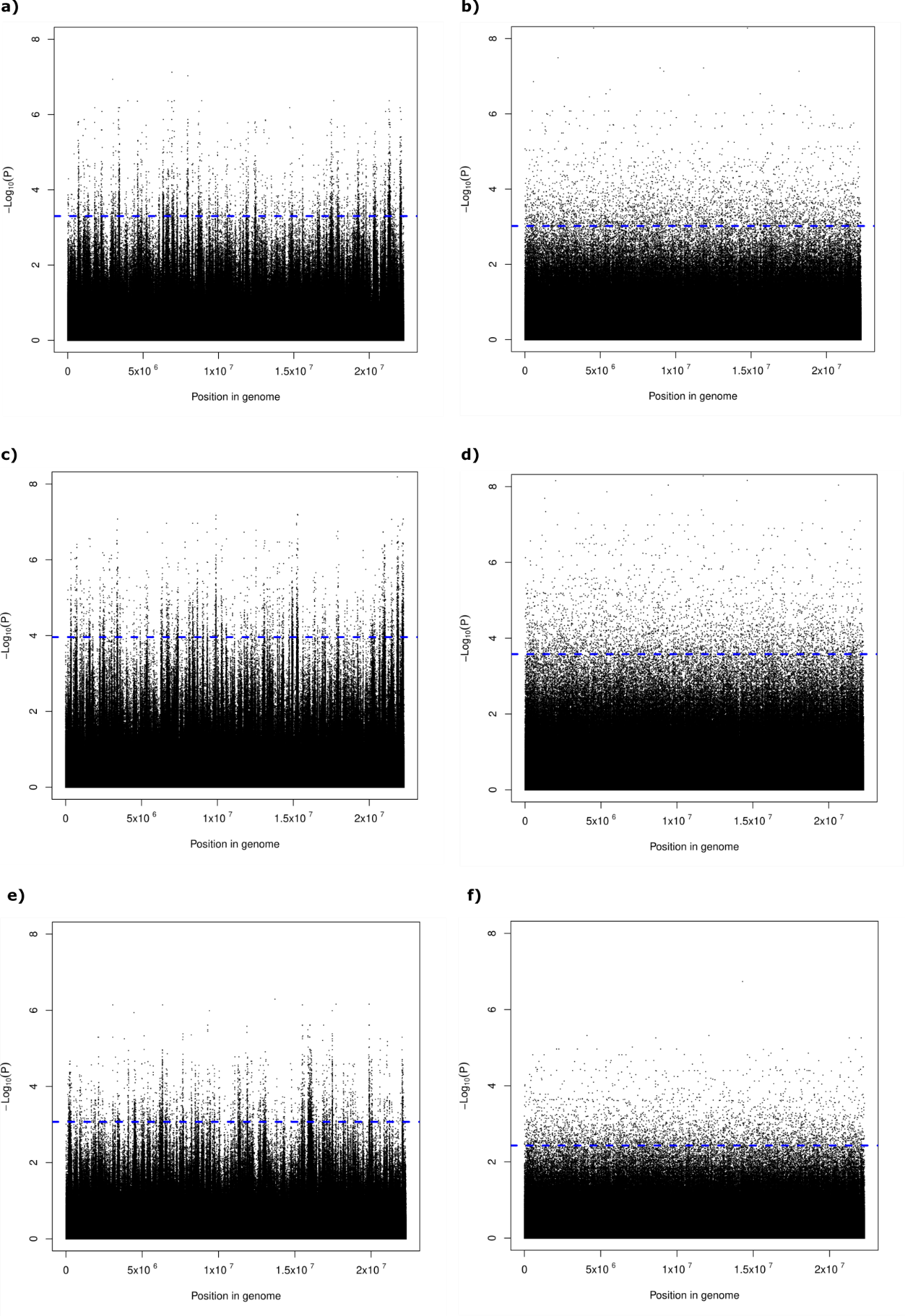

Figure 2: Manhattan plots for the CTRL replicates. Observed results for a) replicate A, c) replicate B and e) replicate C. Simulation for expected drift was done individually b) replicate A, d) replicate B and f) replicate C. Results are from false discovery rate (BH) corrected -Log_10_ probability values from chi-square test. The dashed blue horizontal line indicates the chosen significance threshold (significant quantile above 99.90%).

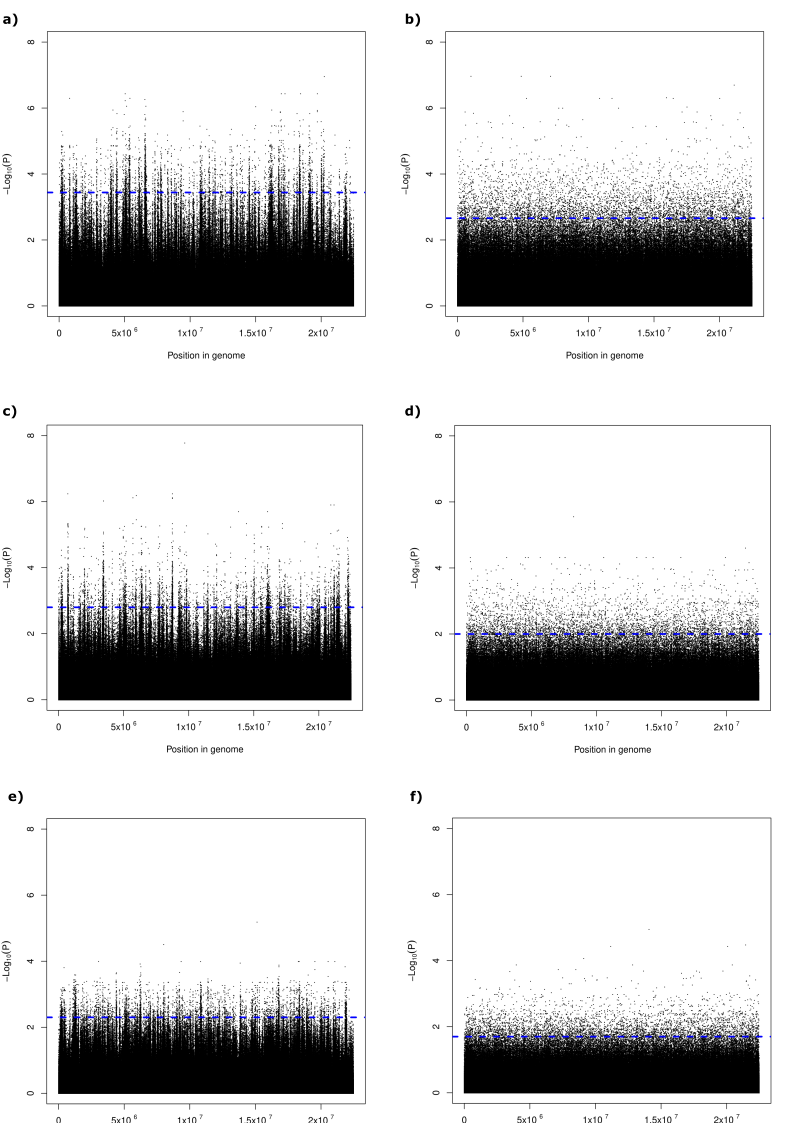

Figure 3: Manhattan plots for the Cd replicates. Observed results for a) replicate A, c) replicate B and e) replicate C. Simulation for expected drift was done individually b) replicate A, d) replicate B and f) replicate C. Results are from false discovery rate (BH) corrected -Log_10_ probability values from chi-square test. The dashed blue horizontal line indicates the chosen significance threshold (significant quantile above 99.90%).

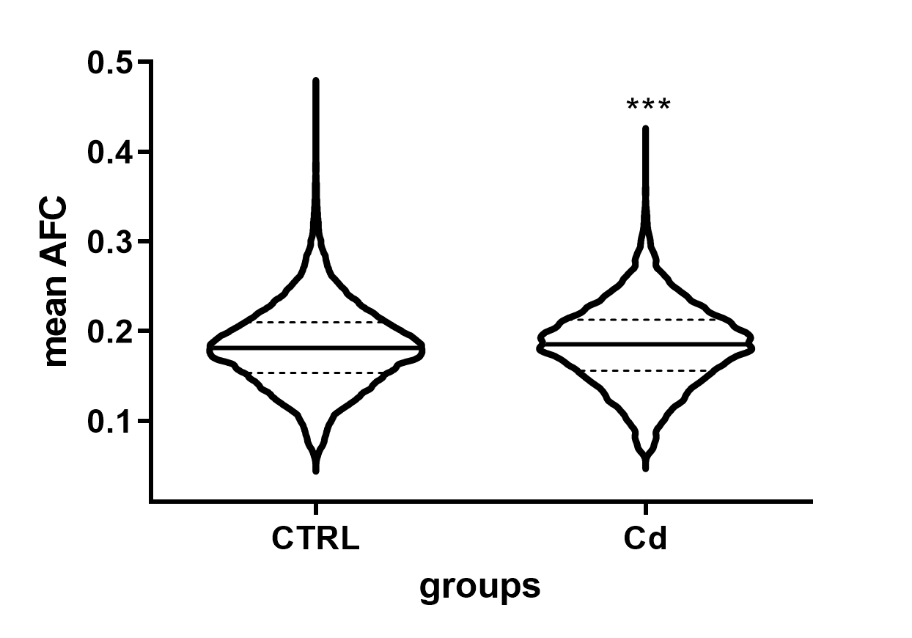

Figure 3: Violin plot of the mean AFCs of the filtered SPNs from the control (CTRL) and Cd pre-treated groups (Cd). Black full line within represents the median, the dashed lines represent the third (up) and first (low) quartiles. Asterisks indicate significant difference between the groups (***p<0.001).

Table 2: List of significantly enriched GO biological functions associated with the genes correlated to the investigated haplotypes and selection targets in control (CTRL) and Cd pre-treated (Cd) groups.

|  | **GO.ID** | **term** | **reference** | **significant** | **p-value** |
| --- | --- | --- | --- | --- | --- |
| **CTRL** | GO:0006355 | regulation of transcription, DNA-template | 378 | 101 | 6.50e-06 |
|  | GO:0007186 | G protein-coupled receptor signaling pathway | 141 | 45 | 0.00059 |
|  | GO:0006813 | potassium ion transport | 45 | 18 | 0.00061 |
|  | GO:0055085 | transmembrane transport | 482 | 118 | 0.00426 |
|  | GO:0006094 | gluconeogenesis | 5 | 4 | 0.00535 |
|  | GO:0007165 | signal transduction | 445 | 126 | 0.00832 |
|  | GO:0007156 | homophilic cell adhesion via plasma membrane | 25 | 10 | 0.01138 |
|  | GO:0034220 | ion transmembrane transport | 108 | 28 | 0.01395 |
|  | GO:0007166 | cell surface receptor signaling pathway | 68 | 24 | 0.0157 |
|  | GO:0051260 | protein homooligomerization | 14 | 7 | 0.02263 |
|  | GO:0006811 | ion transport | 312 | 85 | 0.03402 |
|  | GO:0060828 | regulation of canonical Wnt signaling pathway | 3 | 3 | 0.03549 |
|  | GO:0051016 | barbed-end actin filament capping | 2 | 2 | 0.03555 |
|  | GO:0042908 | xenobiotic transport | 2 | 2 | 0.03555 |
|  | GO:0006368 | transcription elongation from RNA polymerase | 7 | 4 | 0.04938 |
|  | GO:0007076 | mitotic chromosome condensation | 5 | 3 | 0.04947 |
| **Cd** | GO:0007608 | sensory perception of smell | 12 | 6 | 0.00048 |
|  | GO:0006355 | regulation of transcription, DNA-template | 378 | 50 | 0.00448 |
|  | GO:0007186 | G protein-coupled receptor signaling pathway | 141 | 25 | 0.0072 |
|  | GO:0005977 | glycogen metabolic process | 7 | 3 | 0.00961 |
|  | GO:0070588 | calcium ion transmembrane transport | 20 | 6 | 0.01019 |
|  | GO:0055114 | oxidation-reduction process | 438 | 57 | 0.01944 |
|  | GO:0007218 | neuropeptide signaling pathway | 3 | 2 | 0.02702 |
|  | GO:0008272 | sulfate transport | 8 | 3 | 0.03624 |
|  | GO:0035556 | intracellular signal transduction | 160 | 24 | 0.04096 |

GO terms were obtained using topGO with the wheight01 algorithm.

**S4 - Gene expression analysis**

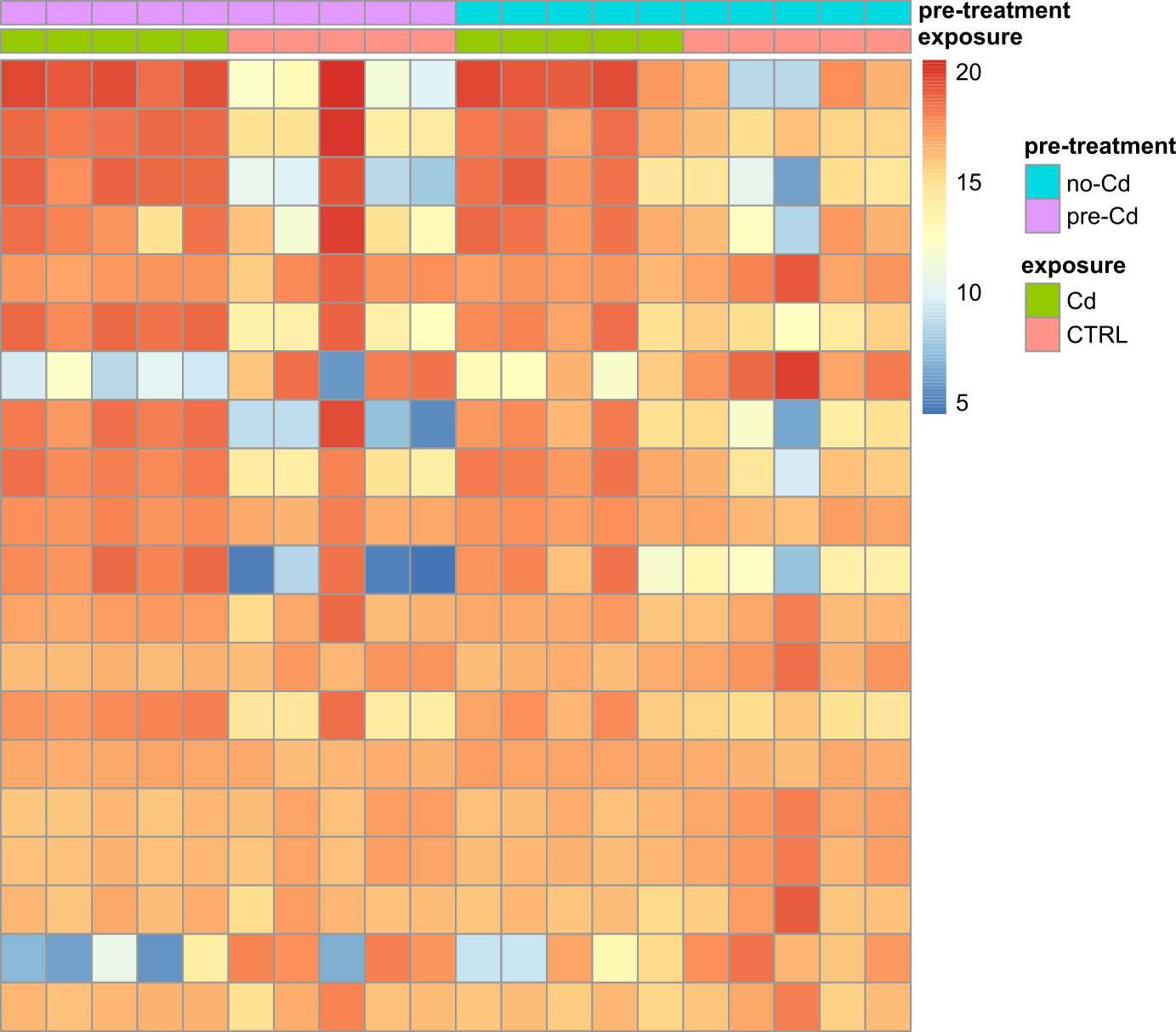

Figure 1: Heatmap showing the expression patterns of genes among the different samples. Pre-treatment is showed as no-Cd (blue) or pre-Cd (purple) and exposure condition is represented by CTRL (pink) or Cd (green). A total of four groups each consisting of five replicates are showed.

Table 1: List of significantly enriched GO biological functions associated with the differentially expressed genes in the Cd pre-treatment and Cd exposure conditions.

|  | **GO.ID** | **term** | **reference** | **significant** | **p-value** |
| --- | --- | --- | --- | --- | --- |
| **pre-treatment** | GO:0007017 | microtubule-based process | 89 | 11 | 0.0037 |
|  | GO:0007018 | microtubule-based movement | 48 | 6 | 0.0258 |
|  | GO:1902476 | chloride transmembrane transport | 1 | 1 | 0.0268 |
|  | GO:0071477 | cellular hypotonic salinity response | 1 | 1 | 0.0268 |
|  | GO:0060294 | cilium movement involved in cell motility | 1 | 1 | 0.0268 |
|  | GO:0018293 | protein-FAD linkage | 1 | 1 | 0.0268 |
|  | GO:0042554 | superoxide anion generation | 1 | 1 | 0.0268 |
| **Cd exposure** | GO:0015986 | ATP synthesis coupled proton transport | 12 | 12 | 1.50e-08 |
|  | GO:0055114 | oxidation-reduction process | 400 | 145 | 6.80e-07 |
|  | GO:0006099 | tricarboxylic acid cycle | 15 | 10 | 0.0003 |
|  | GO:0007601 | visual perception | 11 | 8 | 0.00053 |
|  | GO:0007602 | phototransduction | 11 | 8 | 0.00053 |
|  | GO:0006120 | mitochondrial electron transport (NADH) | 5 | 5 | 0.00056 |
|  | GO:0006122 | mitochondrial electron transport (citC) | 3 | 3 | 0.01125 |
|  | GO:0015671 | oxygen transport | 16 | 8 | 0.01408 |
|  | GO:0019752 | carboxylic acid metabolic process | 160 | 54 | 0.01597 |
|  | GO:0006465 | signal peptide processing | 6 | 4 | 0.02551 |
|  | GO:0055085 | transmembrane transport | 450 | 128 | 0.03232 |
|  | GO:0006777 | Mo-molybdopterin cofactor biosynthetic process | 4 | 3 | 0.03745 |
|  | GO:0090481 | pyrimidine nucleotide-sugar transmembrane transport | 4 | 3 | 0.03745 |
|  | GO:0051276 | chromosome organization | 102 | 24 | 0.04928 |
|  | GO:0022900 | electron transport chain | 13 | 12 | 0.04951 |
|  | GO:0046488 | phosphatidylinositol metabolic process | 31 | 12 | 0.0499 |

Number of genes associated with the term in the reference transcriptome (reference) and in the respective DEG lists (significant). GO terms were obtained using topGO with the wheight01 algorithm. Some significant terms are only represented by a single contig, which may affect any interpretation drawn from them.
